# Systematic analysis of the *Myxococcus xanthus* developmental gene regulatory network supports posttranslational regulation of FruA by C-signaling

**DOI:** 10.1101/415331

**Authors:** Shreya Saha, Pintu Patra, Oleg Igoshin, Lee Kroos

## Abstract

Upon starvation *Myxococcus xanthus* undergoes multicellular development. Rod-shaped cells move into mounds in which some cells differentiate into spores. Cells begin committing to sporulation at 24-30 h poststarvation, but the mechanisms governing commitment are unknown. FruA and MrpC are transcription factors that are necessary for commitment. They bind cooperatively to promoter regions and activate developmental gene transcription, including that of the *dev* operon. Leading up to and during the commitment period, *dev* mRNA increased in wild type, but not in a mutant defective in C-signaling, a short-range signaling interaction between cells that is also necessary for commitment. The C-signaling mutant exhibited ∼20-fold less *dev* mRNA than wild type at 30 h poststarvation, despite a similar level of MrpC and only twofold less FruA. Boosting the FruA level twofold in the C-signaling mutant had little effect on the *dev* mRNA level, and *dev* mRNA was not less stable in the C-signaling mutant. Neither did high cooperativity of MrpC and FruA binding upstream of the *dev* promoter explain the data. Rather, our systematic experimental and computational analyses support a model in which C-signaling activates FruA at least ninefold posttranslationally in order to commit a cell to spore formation.

## Introduction

Differentiated cell types are a hallmark of multicellular organisms. Understanding how pluripotent cells become restricted to particular cell fates is a fascinating question and a fundamental challenge in biology. In general, the answer involves a complex interplay between signals and gene regulation. This is true both during development of multicellular eukaryotes (Davidson & Levine, 2008, Frum & Ralston, 2015, Drapek *et al*., 2017) and during transitions in microbial communities that lead to different cell types (van Gestel *et al*., 2015, Norman *et al*., 2015, Bush *et al*., 2015, Kroos, 2017). Bacterial cells in microbial communities adopt different fates as gene regulatory networks (GRNs) respond to a variety of signals, including some generated by other cells. Moreover, we now understand that microbial communities or microbiomes profoundly impact eukaryotic organisms, and vice versa (Barratt *et al*., 2017, Jansson & Hofmockel, 2018). Yet the daunting complexity of microbiomes and multicellular eukaryotes impedes efforts to fully understand their interactions in molecular detail. By studying simpler model systems, paradigms can be discovered that can guide investigations of more complex interactions.

A relatively simple model system is provided by the bacterium *Myxococcus xanthus*, which undergoes starvation-induced multicellular development (Yang & Higgs, 2014). In response to starvation, cells generate intracellular and extracellular signals that regulate gene expression (Bretl & Kirby, 2016, Kroos, 2017). The rod-shaped cells alter their movements so that thousands form a mound. Within a mound, cells differentiate into ovoid spores that resist stress and remain dormant until nutrients reappear. The spore-filled mound is called a fruiting body. Other cells adopt a different fate and remain outside the fruiting body as peripheral rods (O’Connor & Zusman, 1991). A large proportion of the cells lyse during the developmental process (Lee *et al*., 2012). What determines whether a given cell in the population forms a spore, remains as a peripheral rod, or undergoes lysis? *M. xanthus* provides an attractive model system to discover how signaling between cells affects a GRN and determines cell fate. Here, we focus on a circuit that regulates commitment to sporulation.

In a recent study, cells committed to spore formation primarily between 24 and 30 h poststarvation (PS), because addition of nutrients to the starving population prior to 24 h PS blocked subsequent sporulation, addition at 24 h PS allowed a few spores to form subsequently, and addition at 30 h PS allowed about tenfold more spores to form (Rajagopalan & Kroos, 2014). At the molecular level, addition of nutrients before or during the commitment period caused rapid proteolysis of MrpC (Rajagopalan & Kroos, 2014), a transcription factor required for fruiting body formation (Sun & Shi, 2001b, Sun & Shi, 2001a).

MrpC appears to directly regulate more than one hundred genes involved in development (Robinson *et al*., 2014), and one well-characterized MrpC target gene, *fruA* (Ueki & Inouye, 2003), codes for another transcription factor required for fruiting body formation (Ogawa *et al*., 1996). FruA and MrpC bind cooperatively to the promoter regions of many genes, and appear to activate transcription (Campbell *et al*., 2015, Lee *et al*., 2011, Mittal & Kroos, 2009a, Mittal & Kroos, 2009b, Robinson et al., 2014, Son *et al*., 2011). In particular, transcription of the *dev* operon appears to be activated by cooperative binding of the two transcription factors at two sites located upstream of the promoter (Campbell et al., 2015). Because mutations in three genes of the *dev* operon (*devTRS*) strongly impair sporulation (Boysen *et al*., 2002, Thony-Meyer & Kaiser, 1993, Viswanathan *et al*., 2007a), the feed-forward loop involving MrpC and FruA regulation of the *dev* operon is an attractive molecular mechanism to control spore formation (Fig. 1). Recent work revealed that products of the *dev* operon act as a timer for sporulation (Rajagopalan & Kroos, 2017). DevTRS negatively autoregulate expression of DevI, which inhibits sporulation if overproduced, and delays sporulation by about 6 h when produced normally (Rajagopalan & Kroos, 2017, Rajagopalan *et al*., 2015) (Fig. 1).

**Fig. 1.**
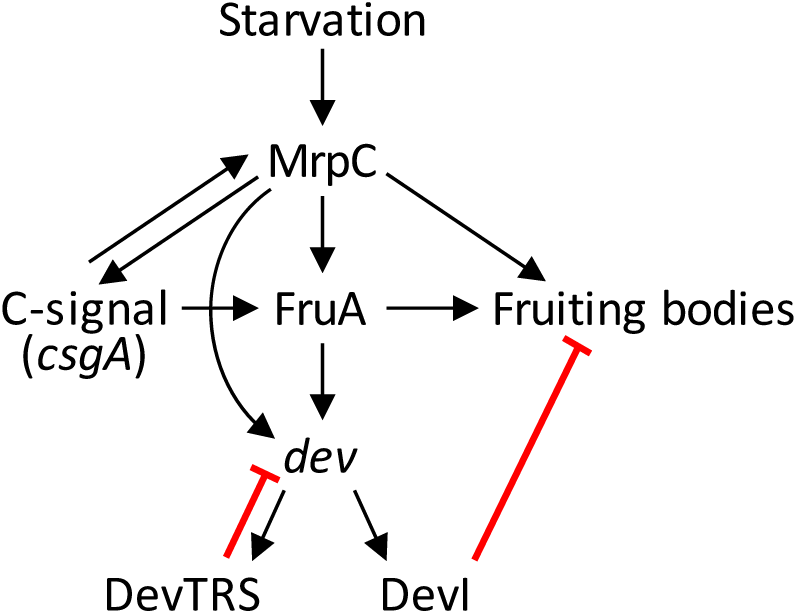
Simplified model of the gene regulatory network governing formation of fruiting bodies. Starvation increases the level of MrpC early in the process (Sun & Shi, 2001b, Sun & Shi, 2001a, Nariya & Inouye, 2006). MrpC causes an increase in C-signal (Sun & Shi, 2001a), the product of *csgA* (Hagen & Shimkets, 1990, Kim & Kaiser, 1990a). MrpC activates transcription of the gene for FruA (Ueki & Inouye, 2003), and C-signal somehow enhances FruA (Ellehauge et al., 1998) and/or MrpC activity (Mittal & Kroos, 2009a). MrpC and FruA bind cooperatively to the promoter region of the *dev* operon and activate transcription (Campbell et al., 2015). The resulting DevTRS proteins negatively autoregulate (Thony-Meyer & Kaiser, 1993, Viswanathan et al., 2007a, Rajagopalan & Kroos, 2017, Rajagopalan et al., 2015). DevI delays spore formation within nascent fruiting bodies (Rajagopalan & Kroos, 2017), but if overproduced, DevI inhibits sporulation (Rajagopalan et al., 2015), which is promoted by MrpC (Sun & Shi, 2001b) and FruA (Ogawa et al., 1996) activity.

Expression of the *dev* operon and many other developmental genes depends on C-signaling (Kroos & Kaiser, 1987), which has been proposed to activate FruA (Ellehauge *et al*., 1998) and/or MrpC (Mittal & Kroos, 2009a) (Fig. 1), although the mechanism of C-signal transduction remains a mystery. Null mutations in the *csgA* gene block C-signaling and sporulation, but the mutants can be rescued by co-development with *csgA+* cells which supply the C-signal (Shimkets *et al*., 1983). C-signaling appears to be a short-range signaling interaction that requires cells to move into alignment (Kim & Kaiser, 1990c, Kim & Kaiser, 1990b, Kroos *et al*., 1988), as they do during mound formation (Sager & Kaiser, 1993). Two theories about the identity of the C-signal have emerged. One theory states that the C-signal is a 17-kDa fragment of CsgA produced by the specific proteolytic activity of PopC at the cell surface (Kim & Kaiser, 1990a, Lobedanz & Sogaard-Andersen, 2003, Rolbetzki *et al*., 2008). The other theory is that diacylglycerols released from the inner membrane by cardiolipin phospholipase activity of intact CsgA are the C-signal (Boynton & Shimkets, 2015). However, in neither case has the signal receptor been identified, so our understanding of C-signaling is incomplete. Likewise, how C-signaling impacts recipient cells is unknown.

One way that C-signaling has been proposed to affect recipient cells is to stimulate autophosphorylation of a histidine protein kinase, which would then transfer the phosphate to FruA (Ellehauge et al., 1998). This model was attractive because FruA is similar to response regulators of two-component signal transduction systems (Ellehauge et al., 1998, Ogawa et al., 1996). Typically, a response regulator is phosphorylated by a histidine protein kinase in response to a signal, thus activating the response regulator to perform a function (Stock *et al*., 2000). The effects of substitutions at the predicted site of phosphorylation in FruA supported the model that FruA is activated by phosphorylation on D59 (Ellehauge et al., 1998). However, a histidine protein kinase capable of phosphorylating FruA has not been identified. Also, several observations suggest that FruA may not be phosphorylated. Most notably, D59 of FruA is present in an atypical receiver domain that lacks a conserved metal-binding residue normally required for phosphorylation to occur, and treatment of FruA with small-molecule phosphodonors did not increase its DNA-binding activity (Mittal & Kroos, 2009a). The receiver domain of FruA was shown to be necessary for cooperative binding with MrpC to DNA, so it was proposed that C-signaling may affect activity of MrpC and/or FruA (Mittal & Kroos, 2009a) (Fig. 1).

The regulation of MrpC has been reported to be complex, involving autoregulation, phosphorylation, proteolytic processing, binding to a toxin protein, and stability (Sun & Shi, 2001b, Nariya & Inouye, 2005, Nariya & Inouye, 2006, Nariya & Inouye, 2008, Schramm *et al*., 2012, Rajagopalan & Kroos, 2014, McLaughlin *et al*., 2018). Also, since MrpC is similar to CRP family transcription factors that bind cyclic nucleotides (Sun & Shi, 2001b), MrpC activity could be modulated by nucleotide binding, so there are many ways in which C-signaling could affect MrpC activity (Mittal & Kroos, 2009a).

Here, using synergistic experimental and computational approaches, we investigate the impact of C-signaling on a circuit that regulates commitment to sporulation by focusing on the feed-forward loop involving MrpC and FruA control of *dev* operon transcription (Fig. 1). We describe methods to systematically and quantitatively study the developmental process. Using these methods we measure the levels of GRN components in wild type and in mutants (e.g., a *csgA* mutant unable to produce C-signal) during the period leading up to and including commitment to spore formation. We then formulate a mathematical model for the steady-state concentration of *dev* mRNA and use the model to computationally predict the magnitude of potential regulatory effects of C-signaling that would be required to explain our data. By testing the predictions, some potential regulatory mechanisms are ruled out and at least ninefold activation of FruA by C-signaling is supported.

## Results

### M. xanthus *development can be studied systematically*

We first established quantitative assays to analyze cellular and molecular changes during *M. xanthus* development. To facilitate collection of sufficient cell numbers for counting, as well as for RNA and protein measurements, development was induced by starvation under submerged culture conditions. Cells adhere to the bottom of a plastic well or dish, and develop under a layer of buffer. Prior to cell harvest, photos were taken to document phenotypic differences between strains. As expected, wild-type strain DK1622 formed mounds by 18 h poststarvation (PS) and the mounds matured into compact, darkened fruiting bodies at 36 to 48 h PS (Fig. 2). In contrast, *csgA* and *fruA* null mutants failed to progress beyond forming loose aggregates. A *devI* null mutant was similar to wild type (WT), whereas a *devS* null mutant formed mounds slowly and they failed to darken. Developing populations were harvested at the times indicated in Figure 2 to measure cellular and molecular changes.

**Fig. 2.**
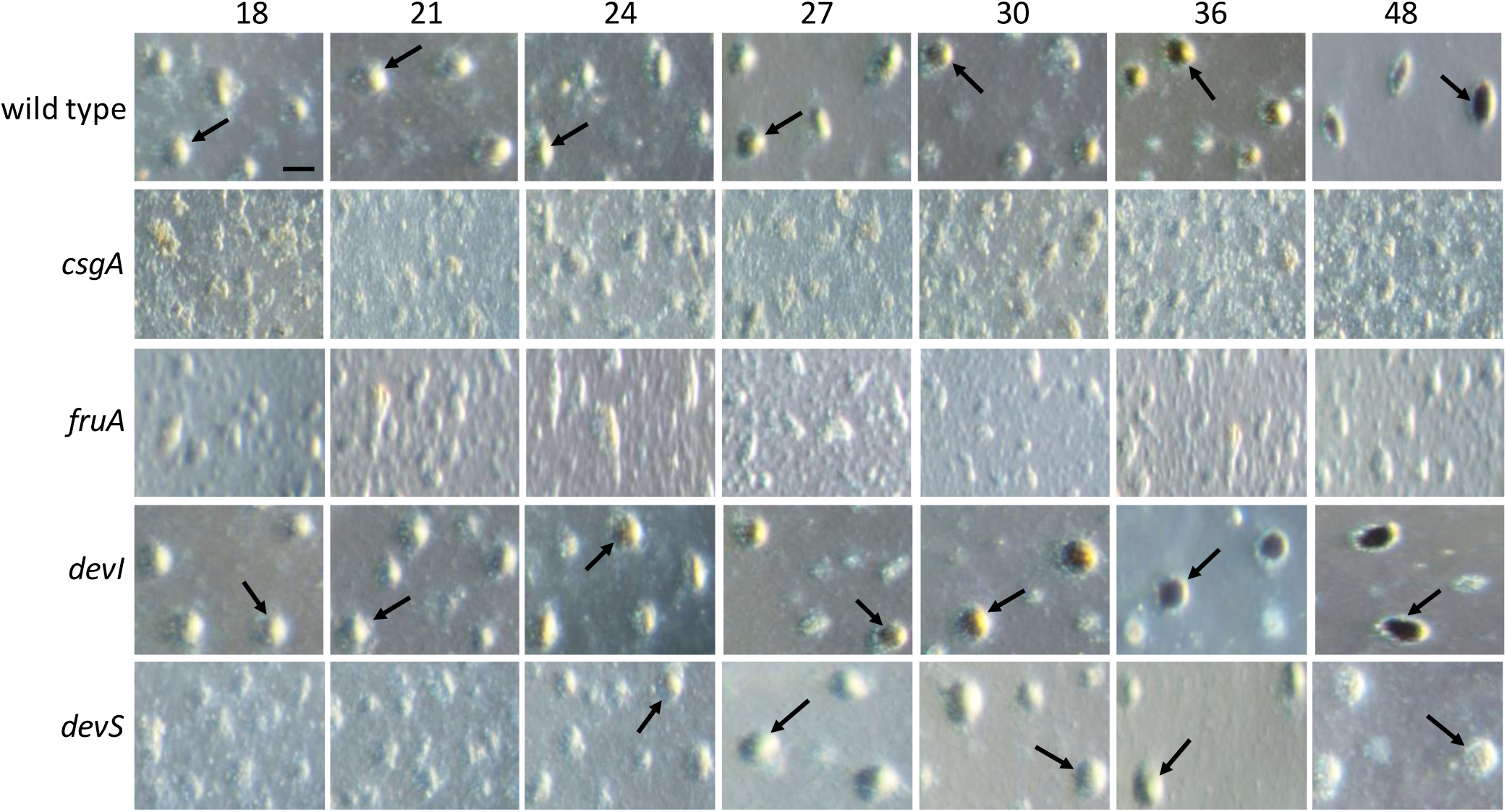
Development of *M. xanthus* strains. Wild-type DK1622 and its indicated mutant derivatives were subjected to starvation under submerged culture conditions and images were obtained at the indicated number of hours poststarvation (PS). DK1622 formed mounds by 18 h PS (an arrow points to one). The *csgA* and *fruA* mutants failed to form mounds, the *devI* mutant was similar to DK1622, and the *devS* mutant formed mounds later, by 24 h PS, but the mounds failed to darken at later times. Bar, 100 μm. Similar results were observed in at least three biological replicates.

To quantify changes at the cellular level, we counted the total number of cells (after fixation and dispersal, so that rod-shaped cells, spores, and cells in transition between the two were counted) and the number of sonication-resistant spores in the developing populations. We also counted the number of rod-shaped cells at the time when development was initiated by starvation (T_0_). By subtracting the number of sonication-resistant spores from the total cell number, we determined the number of sonication-sensitive cells. About 30% of the wild-type cells present at T_0_ remained as sonication-sensitive cells at 18 h PS (Fig. S1A), consistent with the suggestion that the majority of cells lyse early during development under submerged culture conditions, which was based on the decrease in the total protein concentration of developing cultures (Rajagopalan & Kroos, 2014). The number of sonication-sensitive cells continued to decline after 18 h PS, reaching ∼4% of the T_0_ number by 48 h PS (Fig. S1A). Spores were first observed at 27 h PS and the number rose to ∼1% of the T_0_ number by 48 h PS (Fig. S1B). The *devI* mutant was similar to WT, except spores were first observed 6 h earlier at 21 h PS, as reported recently (Rajagopalan & Kroos, 2017). The *csgA, fruA*, and *devS* mutants failed to make a detectable number of spores (at a detection limit of 0.04% of the T_0_ number) and appeared to be slightly delayed relative to WT and the *devI* mutant in terms of the declining number of sonication-sensitive cells (Fig. S1 and Table S1). We conclude that at the cellular level during the time between 18 and 30 h PS (when we measured RNA and protein levels as described below), the developing populations decline from 36 ± 4% to 16 ± 6% of the initial rod number and only 0.5 ± 0.2% (WT, *devI*) or <0.04% (*csgA, fruA, devS*) of the cells form sonication-resistant spores (from which the RNAs and proteins we measured would not be recovered based on control experiments). We stopped collecting samples at 30 PS because thereafter the number of sonication-sensitive cells continues to decline and the spore number continues to rise, making RNA and protein more difficult to recover quantitatively, yet many cells are committed at 30 h PS to make spores by 36 h PS even if nutrients are added (Rajagopalan & Kroos, 2014). Hence, we focused on changes at the molecular level between 18 and 30 h PS, the period leading up to and including the time that many cells commit to spore formation. We acknowledge that the populations also contain cells destined to lyse or become peripheral rods, so the magnitude of molecular changes required to commit a cell to the spore fate may be greater than revealed by our population measurements. However, methods of rapidly separating all three cell types from populations or of analyzing molecular changes in individual cells have not yet been reported.

To measure RNA levels of a large number of samples, we adapted methods described previously (Rajagopalan & Kroos, 2014) to a higher-throughput robotic platform for RT-qPCR analysis. Reproducibility of the analysis was tested among biological replicates and two types of technical replicates as illustrated in Figure S2A, for each RNA to be measured, at 24 h PS, the midpoint of our focal period. No normalization was done in this experiment. Each transcript number was derived from a standard curve of genomic DNA subjected to qPCR. For each RNA, we found that the average transcript number and the standard deviation for three cDNA technical replicates from a single RNA sample, three RNA technical replicates from a single biological replicate, and three biological replicates, was not significantly different (single factor ANOVA, α = 0.05) (Fig. S2B-S2E). These results suggest that biological variation in RNA levels at 24 h PS is comparable to technical variation in preparing RNA and cDNA. In subsequent experiments, we measured RNA for at least three biological replicates and we did not perform RNA or cDNA technical replicates. We also note the high abundance of the *mrpC* transcript (∼10%) relative to 16S rRNA, and the lower relative abundance of the *fruA* (∼1%) and *dev* (∼0.1%) transcripts.

We have typically used 16S rRNA as an internal standard for RT-qPCR analysis during *M. xanthus* development (Rajagopalan & Kroos, 2014). The high abundance of *mrpC* transcript relative to 16S rRNA at 24 h PS (Fig. S2B and S2E) raised the possibility that rRNA decreases relative to total RNA at 18 to 30 h PS. To test this possibility, we measured the 16S rRNA level per μg of total RNA from 18 to 30 h PS. Figure S3A shows that the level does not change significantly (single factor ANOVA, α = 0.05), validating 16S rRNA as an internal standard for subsequent experiments. We also found that the total RNA yield per cell does not change significantly from 18 to 30 h PS (single factor ANOVA, α = 0.05) (Fig. S3B), consistent with the finding that the 16S rRNA level does not change significantly, since the majority of total RNA is rRNA.

To measure protein levels, a portion of each well-mixed developing population was immediately added to sample buffer, boiled, and frozen for subsequent semi-quantitative immunoblot analysis (Rajagopalan & Kroos, 2017). The rest of the population was used for cell counting and RNA analysis as described above and in the Experimental Procedures.

### *Levels of MrpC and FruA fail to account for the low level of* dev *mRNA in a* csgA *mutant*

By systematically quantifying protein and mRNA levels during the period leading up to and including the time that cells commit to spore formation, we investigated whether the GRN shown in Figure 1 could account for observed changes over time in WT and in mutants. In particular, we were interested in whether changes in the levels of MrpC and/or FruA proteins could account for the observed changes in the level of *dev* mRNA, since MrpC and FruA bind cooperatively to the *dev* promoter region and activate transcription (Campbell et al., 2015). In WT, we found that the MrpC level decreased about 1.5-fold on average from 18 to 30 h PS (Fig. 3A) and the FruA level rose about 1.5-fold on average (Fig. 3B), whereas the *dev* mRNA level rose about threefold on average (Fig. 4A). In each case, the fold-change was small and the variation between biological replicates was large, so the result of a single factor ANOVA (α = 0.05) for each time course did not support a significant difference. We reasoned that cooperative binding of MrpC and FruA could easily account for the threefold rise on average in *dev* mRNA. We also measured the levels of *mrpC* and *fruA* mRNA. The *mrpC* mRNA level changed very little on average (Fig. 4B), but the *fruA* mRNA level decreased about twofold on average after 18 h PS (Fig. 4C), in contrast to the 1.5-fold rise on average in the FruA protein level (Fig. 3B), suggesting weak positive posttranscriptional regulation of the FruA level during the period of commitment to spore formation.

**Fig. 3.**
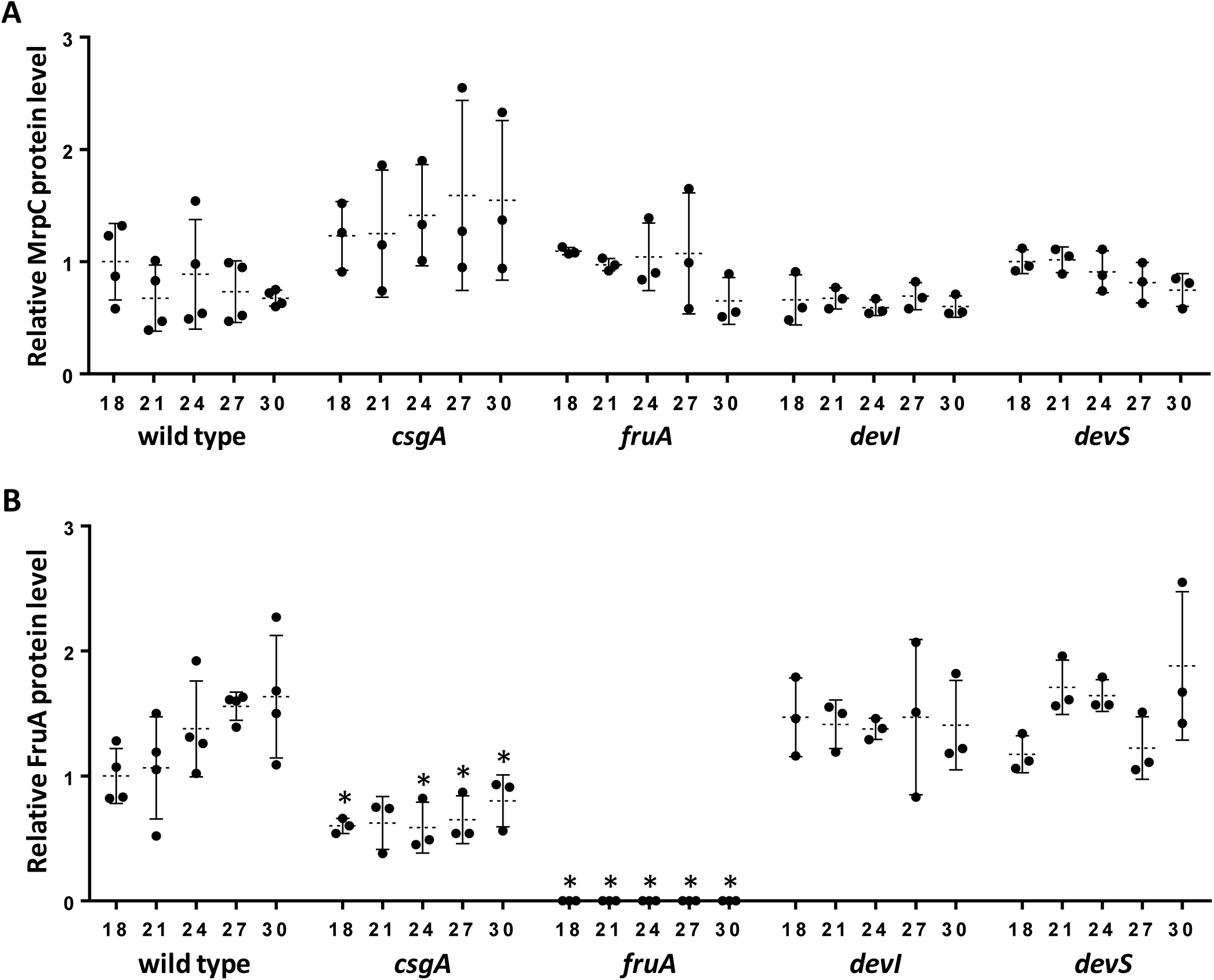
Levels of MrpC and FruA during *M. xanthus* development. Wild-type DK1622 and its indicated mutant derivatives were subjected to starvation under submerged culture conditions and samples were collected at the indicated number of hours poststarvation (PS) for measurement of MrpC (A) and FruA (B) by immunoblot. Graphs show the data points and average of at least three biological replicates, relative to wild-type DK1622 at 18 h PS, and error bars show one standard deviation. Asterisks indicate a significant difference (*p* < 0.05 in Student’s two-tailed *t*-tests) from wild type at the corresponding time PS.

**Fig. 4.**
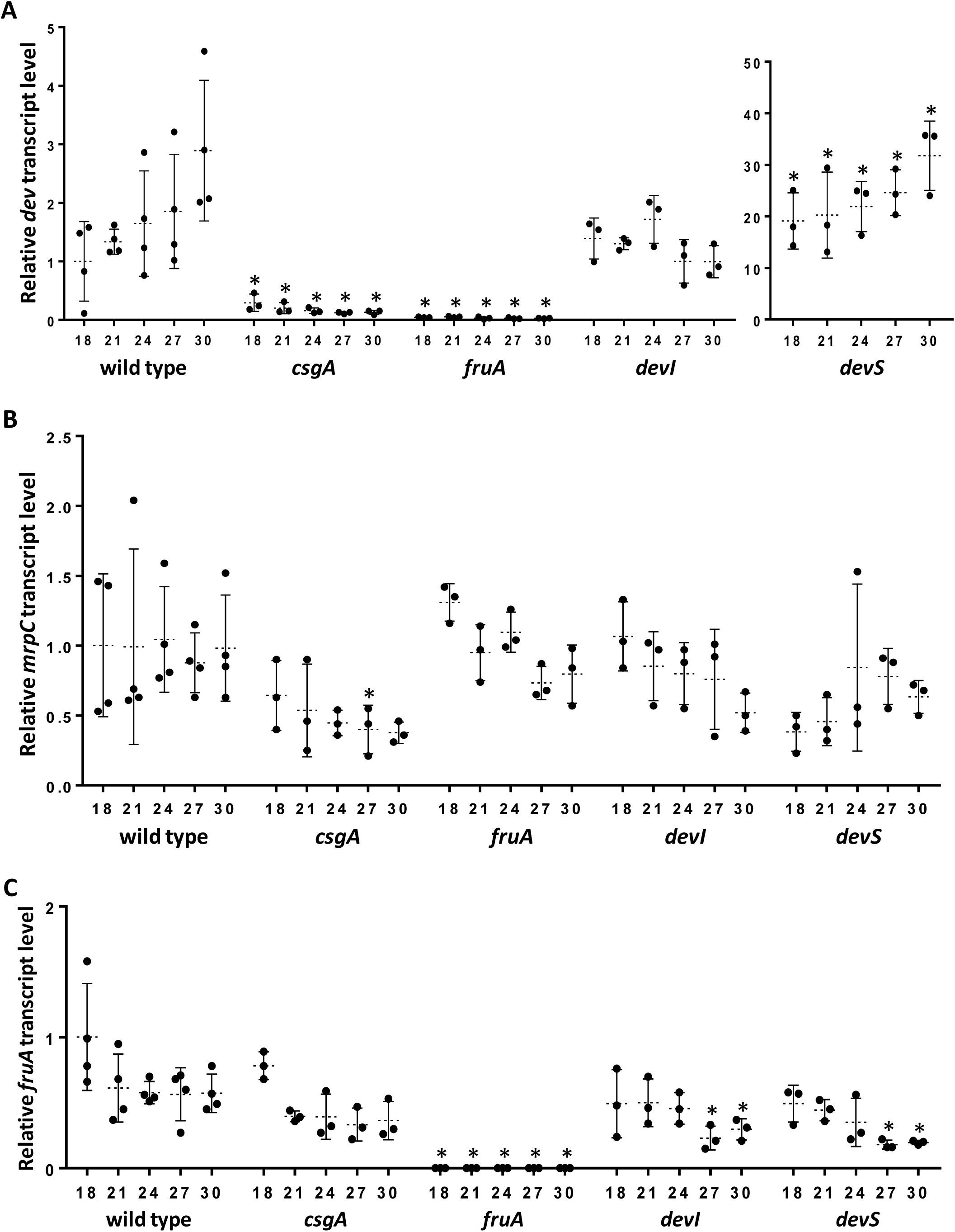
Transcript levels during *M. xanthus* development. Wild-type DK1622 and its indicated mutant derivatives were subjected to starvation under submerged culture conditions and samples were collected at the indicated number of hours poststarvation (PS) for measurement of *dev* (A), *mrpC* (B), and *fruA* (C) transcript levels by RT-qPCR. Graphs show the data points and average of at least three biological replicates, relative to wild-type DK1622 at 18 h PS, and error bars show one standard deviation. Asterisks indicate a significant difference (*p* < 0.05 in Student’s two-tailed *t*-tests) from wild type at the corresponding time PS.

To investigate how C-signaling affects the GRN shown in Figure 1, we measured protein and mRNA levels in the *csgA* null mutant. In agreement with earlier studies suggesting that C-signaling activates FruA (Ellehauge et al., 1998) and/or MrpC (Mittal & Kroos, 2009a), we found very little *dev* mRNA in the *csgA* mutant (Fig. 4A). Notably, the large decrease in the level of *dev* mRNA in the *csgA* mutant compared with WT could not be accounted for by a large decrease in the level of MrpC or FruA. The MrpC level was elevated about 1.5-fold on average in the *csgA* mutant relative to WT at most time points (Fig. 3A), but the differences were not statistically significant (*p* > 0.05 in Student’s two-tailed *t*-tests comparing mutant to WT at each time point). The FruA level was diminished in the *csgA* mutant relative to WT, but only about twofold on average (Fig. 3B). The differences in the FruA level were statistically significant (*p* < 0.05 in Student’s two-tailed *t*-tests) at each time point except 21 h PS (*p* = 0.12), but alone the twofold lower levels of FruA in the *csgA* mutant fail to account for the very low levels of *dev* mRNA.

The *mrpC* and *fruA* mRNA levels were diminished about twofold and 1.5-fold on average, respectively, in the *csgA* mutant relative to WT (Fig. 4B and 4C), but at nearly all time points the differences were not statistically significant (*p* > 0.05 in Student’s two-tailed *t*-tests, except *p* = 0.03 at 27 h for *mrpC* mRNA). The small differences in the level of *fruA* mRNA in the *csgA* mutant relative to WT are especially noteworthy, since they imply that C-signaling has little or no effect on MrpC activity. The results of our *fruA* mRNA measurements agree with published reports using *fruA-lacZ* fusions (Ellehauge et al., 1998, Srinivasan & Kroos, 2004). Furthermore, we found that *fruA* mRNA stability is similar in the *csgA* mutant and in WT at 30 h PS (Fig. S4), indicating that the similar steady-state *fruA* mRNA level we observed (Fig. 4C) reflects a similar rate of synthesis, rather than altered synthesis compensated by altered stability. We conclude that C-signaling does not affect MrpC activity. Therefore, the low level of *dev* mRNA in a *csgA* mutant (Fig. 4A) could be due to failure to activate FruA or to *dev*-specific regulatory mechanisms.

To begin to characterize potential *dev*-specific regulatory mechanisms during the period leading up to and including commitment to sporulation, we measured protein and mRNA levels in the *devS* and *devI* null mutants. The MrpC and FruA levels were similar to WT (Fig. 3). The *dev* mRNA level ranged from 20-fold higher in the *devS* mutant than in WT at 18 h PS, to 10-fold higher at 30 h PS (Fig. 4A), consistent with negative autoregulation by DevS (and DevT and DevR) reported previously (Rajagopalan & Kroos, 2017, Rajagopalan et al., 2015). Unexpectedly, the *dev* mRNA level in the *devI* mutant was about threefold lower than in WT at 30 h PS (Fig. 4A), suggesting that DevI feeds back positively on accumulation of *dev* mRNA, although the difference was not quite statistically significant at the 95% confidence level (*p* = 0.06 in Student’s two-tailed *t*-test). Other differences were that the *fruA* mRNA levels in the *devI* and *devS* mutants were about twofold lower than in WT at 27 and 30 h PS (Fig. 4C), and these were statistically significant (*p* < 0.05 in Student’s two-tailed *t*-tests comparing mutant to WT at each time point). Since the FruA levels in these mutants were similar to those in WT (Fig. 3B), positive posttranscriptional regulation of FruA appeared to occur in the mutants, as well as in WT.

We also measured the *dev* mRNA level in a *ladA* null mutant, because LadA was reported to be a strong positive regulator of *dev* expression based on analysis of transcriptional *lacZ* fusions (Viswanathan *et al*., 2007b). Although the *dev* mRNA was on average twofold lower in the *ladA* mutant than in WT at 18 h PS and the difference was statistically significant (*p* < 0.05 in Student’s two-tailed *t*-tests), significant differences were not observed at 24 or 30 h (Fig. S5A). These results show that LadA does not control the *dev* transcript level during the period (24 to 30 h PS) when cells commit to spore formation under our conditions. Differences in the methods of measuring expression and/or the developmental conditions may account for the different conclusion reached here about the effect of LadA on *dev* expression, as compared with the previous study (Viswanathan et al., 2007b). We note that, as in the previous study, the *ladA* mutant was delayed for mound formation and darkening (Fig. S6A). Formation of sonication-resistant spores was also delayed (Fig. S6B).

To complete our characterization of the GRN shown in Figure 1, we also measured protein and mRNA levels in the *fruA* and *mrpC* null mutants. We did not collect samples of the *mrpC* mutant at as many time points since we expected little or no expression of GRN components. As expected, neither MrpC nor FruA were detected in the *mrpC* mutant (Fig. S7). In the *fruA* mutant, the MrpC level was similar to WT and, as expected, FruA was not detected (Fig. 3). Also as expected, in the *fruA* mutant the *fruA* mRNA was not detected, the *dev* mRNA level was very low, and the *mrpC* mRNA level was similar to WT (Fig. 4). Since the *mrpC* mutant had an in-frame deletion of codons 74 to 229 (Sun & Shi, 2001b), we were able to design primers for RT-qPCR analysis that should detect the shorter *mrpC* transcript. Surprisingly, the *mrpC* mutant exhibited an elevated level of *mrpC* transcript compared with WT at 18 and 24 h PS (Fig. S8A). The result was surprising since expression of an *mrpC-lacZ* fusion had been reported to be abolished in the *mrpC* mutant, which had led to the conclusion that MrpC positively autoregulates (Sun & Shi, 2001b). We considered the possibility that the shorter transcript in the *mrpC* mutant is more stable than the WT transcript, but the transcript half-lives after addition of rifampicin did not differ significantly (Fig. S9). We conclude that MrpC negatively regulates the *mrpC* transcript level. While this work was in progress, McLaughlin *et al*. reached the same conclusion (McLaughlin et al., 2018). In all other respects, the *mrpC* mutant yielded expected results. The *fruA* and *dev* transcripts were very low (Fig. S8B and S8C), consistent with the expectations that MrpC is required to activate *fruA* transcription (Ueki & Inouye, 2003) and that MrpC and FruA are required to activate *dev* transcription (Campbell et al., 2015, Ellehauge et al., 1998, Viswanathan et al., 2007b). Also, the *mrpC* mutant failed to progress beyond forming loose aggregates (Fig. S10), appeared to be slightly delayed relative to WT in terms of the declining number of sonication-sensitive cells (Fig. S11A), and failed to make a detectable number of spores (at a detection limit of 0.04% of the T_0_ number) (Fig. S11B).

Taken together, our systematic, quantitative measurements of components of the GRN shown in Figure 1 imply that failure to activate FruA and/or *dev*-specific regulatory mechanisms may account for the low level of *dev* mRNA in a *csgA* mutant. Given the complex feedback architecture of *dev* regulation (i.e., strong negative feedback by DevTRS and weak positive feedback by DevI at 30 h PS), delineating the effects of C-signaling on the *dev* transcript level requires a mathematical modeling approach.

### *Mathematical modeling suggests several mechanisms that could explain the low level of dev mRNA in the* csgA *mutant*

The observed small differences in the levels of MrpC and FruA in the *csgA* mutant relative to WT do not account for the very low level of *dev* mRNA in the *csgA* mutant. To evaluate plausible mechanisms that may explain these experimental findings, we quantitatively analyzed transcriptional regulation of *dev* by formulating a mathematical model that expresses the *dev* mRNA concentration as a function of the regulators MrpC, FruA, DevI, and DevS. MrpC and FruA bind cooperatively to the *dev* promoter region and activate transcription (Campbell et al., 2015). Our results suggest that DevI is a weak positive regulator and DevS is a strong negative regulator of *dev* transcription by 30 h PS (Fig. 4A).

Incorporating these effects into a transcriptional regulation model, we express the concentration of *dev* mRNA as a product of three regulation functions (П_FM_, П_I_, П_S_) divided by the transcript degradation rate *δ*_*dev*_ (see Experimental Procedures for detailed explanation):

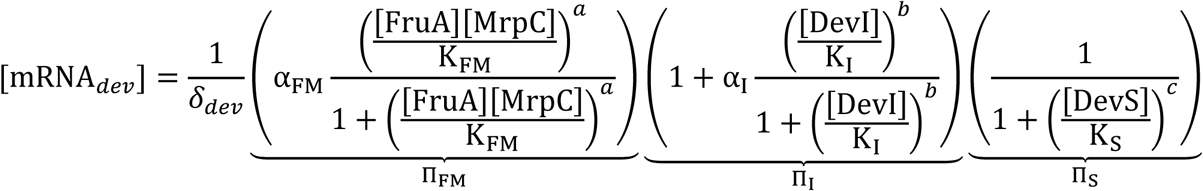

Here, we use a quasi-steady state approximation for the mRNA levels by taking advantage of the fact that mRNA decay (with half-lives typically in minutes) is much faster than our experimental measurement times (in hours). This allows us to assume a rapid equilibrium between the rate of *dev* transcription and the decay of its mRNA, which leads to the above equation, in which α_FM_, α_I_, δ_*dev*_, a, b, c, K_FM_, K_I_ and K_S_ are parameters characterizing promoter regulation. We assume that these biochemical parameters are not a function of the genetic background and, therefore, in the strains in which *dev* mRNA was measured (e.g., the *csgA* mutant), the concentration of *dev* mRNA is determined by the concentrations of proteins (indicated by square brackets in the equation), more specifically the concentrations of their transcriptionally active forms (in case there is a posttranslational regulation). To estimate how the different regulation parameters (such as transcription rate, degradation rate, cooperativity constant, etc.) affect the *dev* mRNA level, we first constrain the model parameters by the experimental result shown in Figure 3B, [FruA]_WT_/ [FruA]_*cagA*_ ≅ 2, and search for parameters that can result in the observed 22-fold difference in [MRNA_*dev*_] in WT relative to the *csgA* mutant at 30 h PS (Fig. 4A).

To estimate the contribution of autoregulation by Dev proteins to their own transcription (i.e., the terms П_I_, П_S_) in WT and the *csgA* mutant, we employ the data from the *devI* and *devS* mutants (Fig. 4A). Specifically, we take the ratio of the *dev* mRNA level in WT to that in *devI* and *devS* mutants to estimate the feedback regulation from DevI and DevS, respectively (see Experimental Procedures for details). We find the contribution from DevI and DevS feedback regulation in WT to be П_I,WT_ = 2.9 and П_S,WT_ = 0.091, respectively. Using these values, we find the contribution from FruA and MrpC regulation to be П_FM,WT_/*δ*_*dev*,WT_ = 11. In the *csgA* mutant, since the *dev* mRNA level is very low, we assume the DevI and DevS protein levels to be low. This gives the contribution of different regulation functions as П_I,*cagA*_ ≈ 1, П_S,*cagA*_ ≈ 1, and П_FM,*cagA*_/*δ*_*dev,cagA*_ = 0.13. In summary, this analysis reveals that the twofold reduction of FruA protein observed in the *csgA* mutant (Fig. 3B) leads to a change of (П_FM,WT_/П_FM,*cagA*_)(*δ*_*dev,cagA*_/*δ*_*dev*,WT_) ≈ 84-fold in the FruA-and MrpC-dependent transcript regulation term. We reasoned that the observed 22-fold reduction in *dev* transcript in the *csgA* mutant relative to WT at 30 h PS (Fig. 4A) could result from a reduction in the FruA-and MrpC-dependent activation rate П_FM_ and/or an increase in the transcript degradation rate *δ*_*dev*_. In what follows we use the mathematical model to predict the magnitude of these effects that would be necessary to explain the observed 22-fold difference in [MRNA_*dev*_].

#### Hypothesis 1: Increase in *dev* transcript degradation rate in the *csgA* mutant

First, we estimate the difference in *dev* transcript degradation rate necessary to explain the observed difference in transcript level between WT and the *csgA* mutant. For this, we make two assumptions. First, we assume that MrpC and FruA bind to the *dev* promoter region with a Hill cooperativity coefficient a = 2 (i.e., the maximum for a single cooperative binding site). Second, we assume that the observed twofold difference in FruA protein level results in a twofold difference in transcriptionally active FruA. Under these assumptions, we vary the remaining unknown parameters to compute the required fold difference in transcript degradation rate for different values of promoter saturation. Our results plotted in Figure 5A show that at least a 20-fold difference in transcript degradation rate is required to explain the transcript data. This experimentally testable prediction will be assessed in a subsequent section. If the results are inconsistent with this prediction, we must conclude that at least one of the two assumptions above is invalid, resulting in the following two alternative hypotheses: the Hill coefficient of MrpC and FruA binding to the *dev* promoter region is much higher than a = 2 and/or the amount of transcriptionally active FruA does not scale with the measured FruA protein level (e.g., if *csgA*-dependent C-signaling is also involved in posttranslational activation of FruA).

**Fig. 5.**
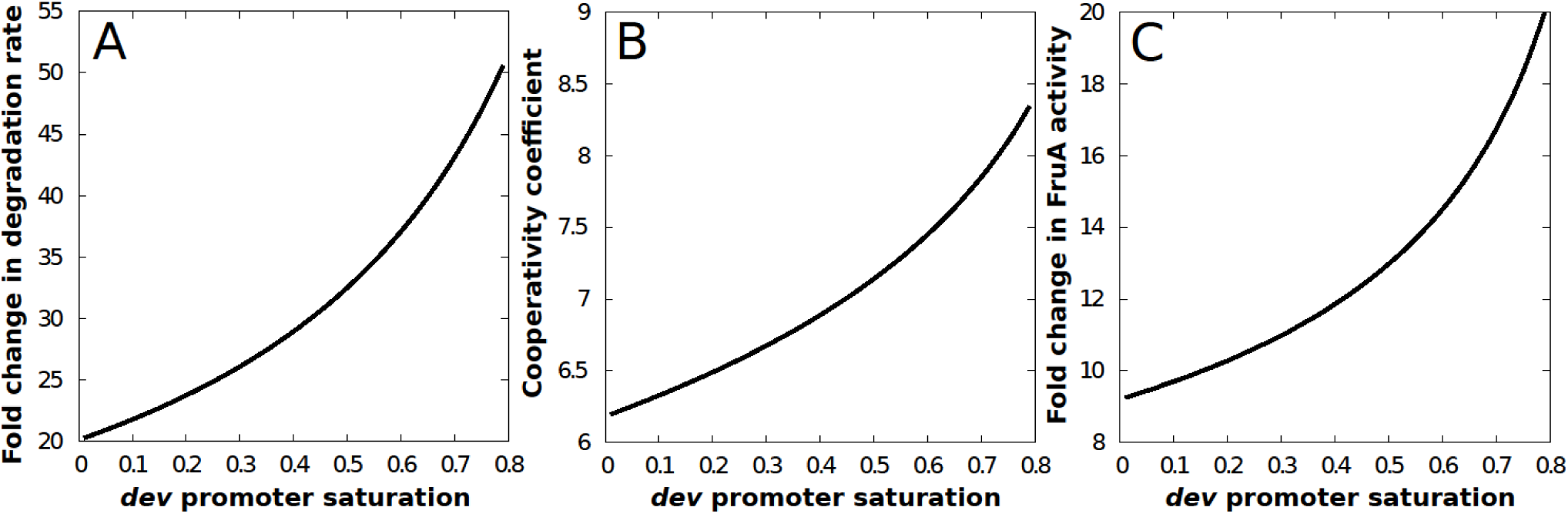
Mathematical modeling of different hypotheses to explain the low *dev* transcript level in a *csgA* mutant. Plots showing the required fold change in *dev* transcript degradation rate in the *csgA* mutant in comparison to wild type (A), cooperativity coefficient for MrpC and FruA binding to the *dev* promoter region (B), and reduction in FruA activity in the *csgA* mutant in comparison to wild type (C), to explain the experimental data for different values of promoter saturation.

#### Hypothesis 2: High cooperativity of MrpC and FruA binding to the *dev* promoter region

Next, we test if a higher binding cooperativity can explain the difference in *dev* transcript level between WT and the *csgA* mutant. We compute the required cooperativity coefficient by assuming the degradation rate does not change between the two strains. Our results plotted in Figure 5B show that the minimum cooperativity coefficient required to explain the experimental results is six for low promoter saturation. In biologically realistic conditions, where promoter saturation is higher; the required cooperativity is even higher. Such a large cooperativity can only be explained if there is more than one site in the promoter region where MrpC and FruA bind with high cooperativity. We know that the *dev* promoter region has at least two MrpC and FruA cooperative binding sites; one is proximal upstream, whereas the other is distal upstream (Campbell et al., 2015). The distal upstream binding site appeared to boost *dev* promoter activity after 24 h PS, based on β-galactosidase activity from a *lacZ* reporter. Hence, in a subsequent section, we study the impact of a distal site deletion on different transcripts (*mrpC, fruA, dev*) and proteins (MrpC, FruA) to test if presence of the distal site contributes to higher cooperativity. If the results are not consistent with the model predictions, we must conclude that the fold difference in active FruA exceeds that observed for the total concentration of each protein (i.e., *csgA*-dependent C-signaling is involved in posttranslational activation of FruA).

#### Hypothesis 3: Posttranslational regulation of FruA activity

To assess the difference in active FruA level required to explain the observed difference in *dev* transcript level, in the absence of other effects, we fix the cooperativity coefficient at a = 2 and assume the transcript degradation rate to be unchanged between WT and the *csgA* mutant. We then use our model to compute the fold difference in active FruA required to achieve a 22-fold reduction in *dev* transcript in the *csgA* mutant relative to WT. Our results plotted in Figure 5C show that at least a ninefold reduction in active FruA is needed in the *csgA* mutant. The reduction in active FruA in the *csgA* mutant would presumably be due to the absence of C-signal-dependent posttranslational activation of FruA, not due to the twofold lower level of FruA protein we observed in the *csgA* mutant relative to WT (Fig. 3B). The reduction in active FruA may be considerably greater than ninefold if the *dev* promoter region approaches saturation (e.g., 20-fold at 80% saturation in Fig. 5C). Also, mathematical modeling of our data at each time point from 18 to 30 h PS yields a similar result (Fig. S12), suggesting that in WT, FruA has already been activated by C-signaling at least ninefold by 18 h PS, and perhaps as much as 30-fold if the *dev* promoter region approaches saturation (righthand panel in Fig. S12).

### *Stability of the* dev *transcript is unchanged in a* csgA *mutant*

To measure the *dev* transcript degradation rate in WT and the *csgA* mutant, we compared the *dev* transcript levels after addition of rifampicin to block transcription at 30 h PS. The average half-life of the *dev* transcript in three biological replicates was 11 ± 6 min in WT and 7 ± 1 min in the *csgA* mutant (Fig. 6), which is not a statistically significant difference (*p* = 0.36 in a Student’s two-tailed *t*-test). We conclude that elevated turnover does not account for the low level of *dev* transcript in the *csgA* mutant. These results allow us to rule out Hypothesis 1.

**Fig. 6.**
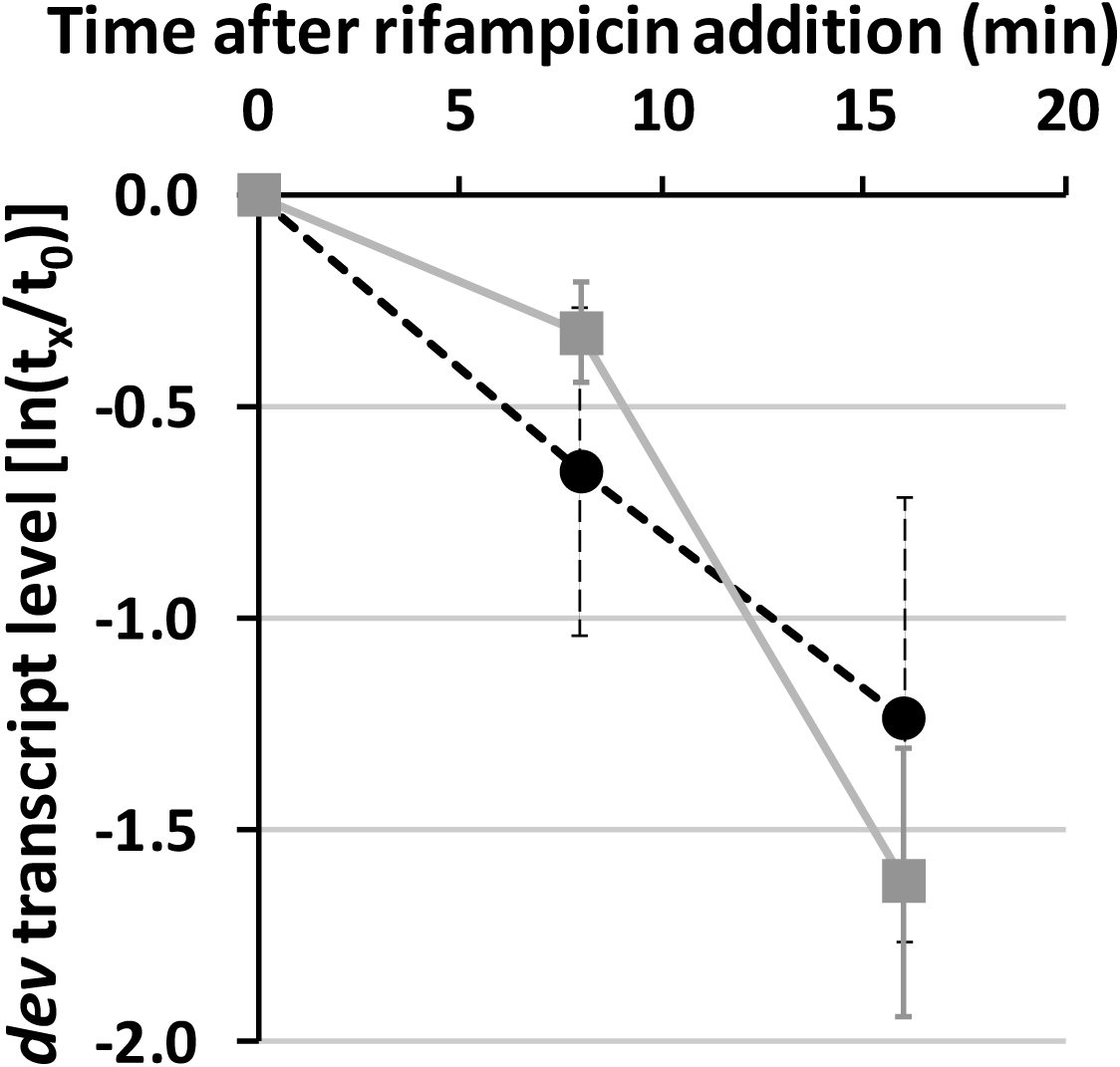
*dev* transcript stability. Wild-type DK1622 and the *csgA* mutant were subjected to starvation under submerged culture conditions for 30 h. The overlay was replaced with fresh starvation buffer containing rifampicin (50 μg/ml) and samples were collected immediately (t_0_) and at the times indicated (t_x_) for measurement of the *dev* transcript level by RT-qPCR. Transcript levels at t_x_ were normalized to that at t_0_ for each of three biological replicates and used to determine the transcript half-life for each replicate. The average half-life and one standard deviation are reported in the text. The graph shows the average ln(t_x_/ t_0_) and one standard deviation for the three biological replicates of wild type (black dashed line) and the *csgA* mutant (gray solid line).

### *The distal upstream binding site for MrpC and FruA has little impact on the* dev *transcript level*

In a previous study, weak cooperative binding of MrpC and FruA to a site located between positions −254 and −229 upstream of the *dev* promoter appeared to boost β-galactosidase activity from a *lacZ* transcriptional fusion about twofold between 24 and 30 h PS, but deletion of the distal upstream site did not impair spore formation (Campbell et al., 2015). These findings suggested that the distal site has a modest impact on *dev* transcription that is inconsequential for sporulation. However, β-galactosidase activity from *lacZ* fused to *dev* promoter segments with different amounts of upstream DNA and integrated ectopically may not accurately reflect the contribution of the distal site to the *dev* transcript level. Therefore, we measured the *dev* transcript level in a mutant lacking the distal site (i.e., DNA between positions −254 and −228 was deleted from the *M. xanthus* chromosome). The level of *dev* transcript in the distal site mutant was similar to WT measured in the same experiment, in this case increasing about twofold from 18 to 30 h PS (Fig. 7). Likewise, the levels of *mrpC* and *fruA* transcripts (Fig. S8) and the corresponding proteins (Fig. S7) were similar in the distal site mutant as compared with WT (*p* > 0.05 in Student’s two-tailed *t*-tests comparing mutant to WT at each time point, except *p* = 0.02 for *fruA* transcript levels at 18 h, *p* = 0.03 for MrpC protein levels at 30 h, and *p* = 0.03 for FruA protein levels at 18 h). The distal site mutant formed mounds by 18 h PS, which matured into compact, darkened fruiting bodies at later times, similar to WT (Fig. S10), and the percentages of sonication-sensitive cells and sonication-resistant spores observed for the distal site mutant were similar to WT (Fig. S11). We conclude that the distal site has little or no impact on the developmental process. In particular, the distal site does not contribute to high cooperativity of MrpC and FruA binding to the *dev* promoter region that could explain the higher level of *dev* transcript in WT than in the *csgA* mutant. These results allow us to rule out Hypothesis 2.

**Fig. 7.**
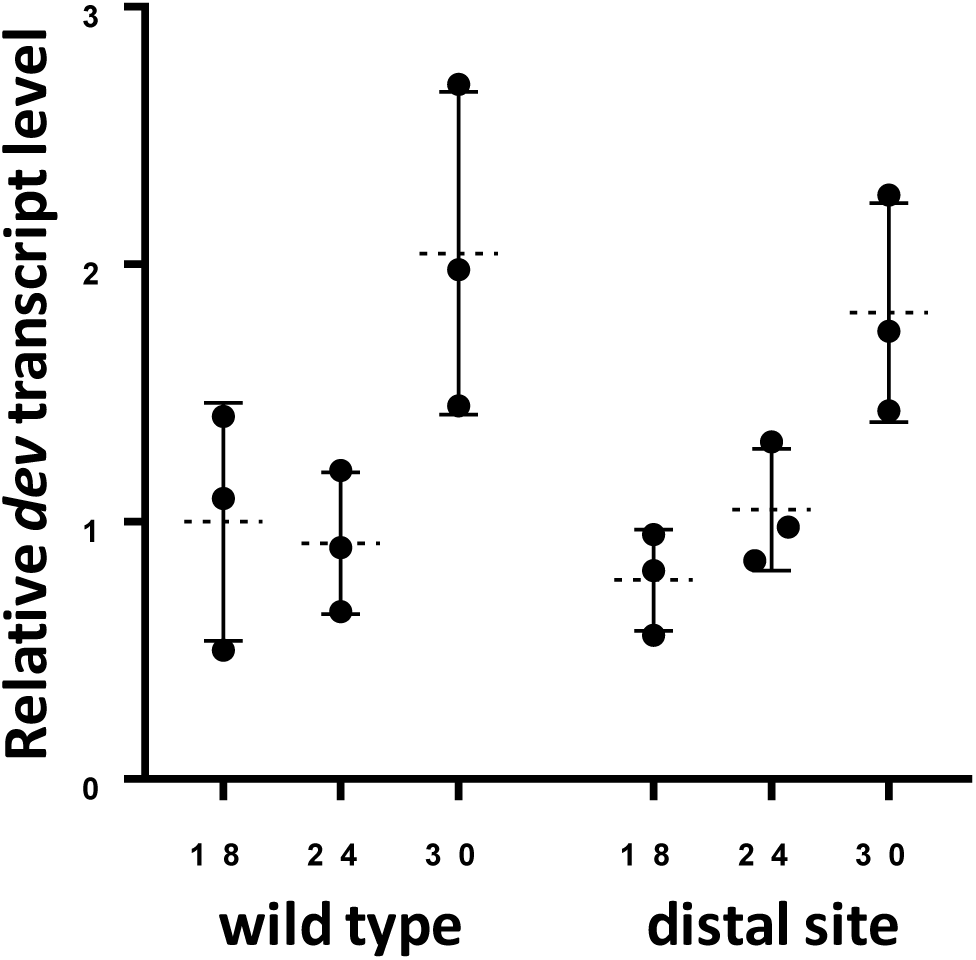
*dev* transcript levels. Wild-type DK1622 and its indicated mutant derivative were subjected to starvation under submerged culture conditions and samples were collected at the indicated number of hours poststarvation (PS) for measurement of *dev* transcript levels by RT-qPCR. Graphs show the data points and average of three biological replicates, relative to wild-type DK1622 at 18 h PS, and error bars show one standard deviation.

### *Boosting the FruA level in the* csgA *mutant has no effect on the* dev *transcript level*

Having ruled out the first two hypotheses, our modeling predicts that the only viable option to explain the effect of the *csgA* null mutation on the *dev* transcript level is Hypothesis 3: at least a ninefold reduction in active FruA is needed in the *csgA* mutant as compared with WT. Specifically, our model showed that the low *dev* transcript level in the *csgA* mutant is not due to its twofold lower FruA level (Fig. 3B), but rather due to a failure to activate FruA in the absence of C-signaling (Fig. 5C and S12). As a result, the model predicts that in the *csgA* mutant most of the FruA remains inactive. To test this prediction, we integrated *fruA* transcriptionally fused to a vanillate-inducible promoter ectopically in the *csgA* mutant. Upon induction the *csgA* P_*van*_-*fruA* strain accumulated a similar level of FruA as WT (Fig. 8A), but the *dev* transcript level remained as low as in the *csgA* mutant (Fig. 8B). Hence, boosting the FruA level in the *csgA* mutant had no effect on the *dev* transcript level, consistent with our prediction and supporting the hypothesis that C-signaling activates FruA at least ninefold.

**Fig. 8.**
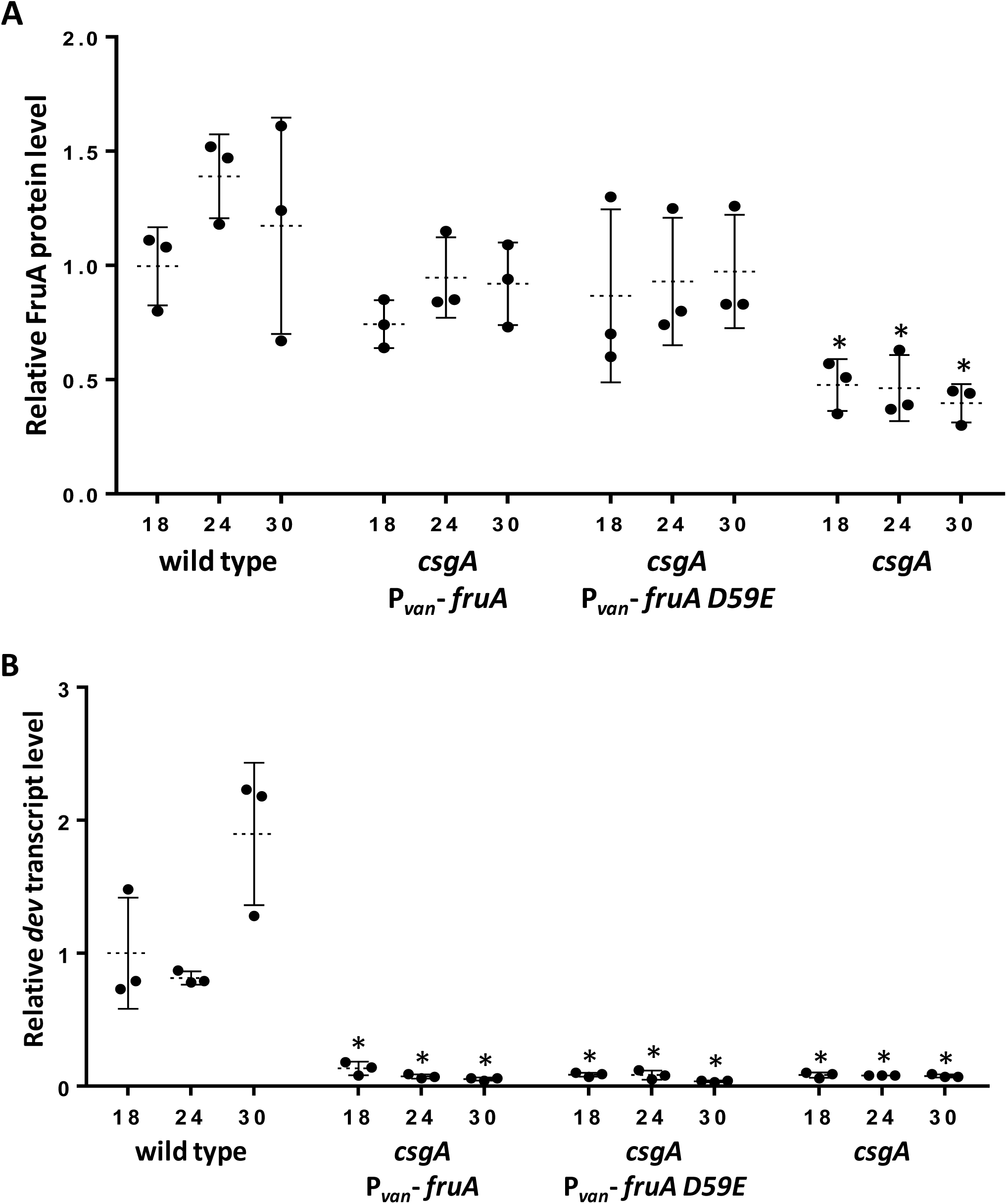
FruA protein and *dev* transcript levels. Wild-type DK1622 and its indicated mutant derivatives were subjected to starvation under submerged culture conditions and samples were collected at the indicated number of hours poststarvation (PS) for measurement of FruA levels by immunoblot (A) and *dev* transcript levels by RT-qPCR (B). Graphs show the data points and average of three biological replicates, relative to wild-type DK1622 at 18 h PS, and error bars show one standard deviation. Asterisks indicate a significant difference (*p* < 0.05 in Student’s two-tailed *t*-tests) from wild type at the corresponding time PS.

Additionally, we tested the previously proposed model that C-signal-dependent phosphorylation of D59 activates FruA (Ellehauge et al., 1998). We created *fruA D59E*, an allele designed to produce FruA D59E with a phosphomimetic substitution in its receiver domain. In some cases, such a substitution causes a response regulator to be constitutively active (Domian *et al*., 1997, Klose *et al*., 1993, Lan & Igo, 1998), including response regulators in the NarL/FixJ subfamily to which FruA belongs (Davlieva *et al*., 2015, Galperin, 2010, Ellehauge et al., 1998). Therefore, if FruA is normally activated by phosphorylation of D59 in response to C-signaling, expression of *fruA D59E* may restore the *dev* transcript level in the *csgA* mutant. We integrated *fruA D59E* transcriptionally fused to a vanillate-inducible promoter ectopically in the *csgA* mutant. Upon induction the *csgA* P_*van*_-*fruA D59E* strain accumulated FruA D59E to a similar level as FruA accumulated by WT (Fig. 8A), but the *dev* transcript level in the *csgA* P_*van*_-*fruA D59E* strain remained as low as in the *csgA* mutant (Fig. 8B). These results suggest that D59 phosphorylation may not be the mechanism by which C-signaling activates FruA.

The *fruA* transcript (Fig. S13A), *mrpC* transcript (Fig. S13B), and MrpC protein (Fig. S14) levels were similar in the *csgA* P_*van*_-*fruA* and *csgA* P_*van*_-*fruA D59E* strains as in WT and the *csgA* mutant. Induction of the *csgA* P_*van*_-*fruA* and *csgA* P_*van*_-*fruA D59E* strains did not rescue development, which failed to progress beyond forming loose aggregates (Fig. S15), and the strains failed to make a detectable number of spores by 48 h PS (Table S1). The induced *csgA* P_*van*_-*fruA* strain appeared to be slightly delayed relative to WT in terms of the declining number of sonication-sensitive cells, like the *csgA* mutant, whereas the induced *csgA* P_*van*_-*fruA D59E* appeared to be defective in attachment, resulting in recovery of less sonication-sensitive cells at 18 h PS (Fig. S16).

As controls, P_*van*_-*fruA* and P_*van*_-*fruA D59E* were integrated ectopically in the *fruA* mutant. Upon induction the *fruA* P_*van*_-*fruA* strain formed mounds by 18 h PS and the mounds matured into compact, darkened fruiting bodies at later times, similar to WT without or with vanillate added (Fig. S17). The induced *fruA* P_*van*_-*fruA D59E* strain formed mounds by 24 h PS, which matured into compact, darkened fruiting bodies, similar to WT. Also, the induced *fruA* P_*van*_-*fruA* and P_*van*_-*fruA D59E* strains exhibited a similar number of sonication-resistant spores as WT at 36 h PS (data not shown). These results show that induction of the *fruA* P_*van*_-*fruA* and *fruA* P_*van*_-*fruA D59E* strains rescued development, presumably because C-signaling activated the FruA and FruA D59E produced, respectively.

### *Other FruA-dependent transcripts are present at very low levels in the* csgA *mutant, even when FruA or FruA D59E is produced*

In addition to the *dev* operon, four other genes or operons appear to be activated by cooperative binding of FruA and MrpC in their promoter region (Lee et al., 2011, Mittal & Kroos, 2009a, Mittal & Kroos, 2009b, Son et al., 2011). These genes, designated *fmg* for “<u>F</u>ruA-and <u>M</u>rp-regulated <u>g</u>ene”, also depend on C-signaling for expression, based on analysis of *lacZ* fusions (Kroos & Kaiser, 1987). We measured the levels of *fmgA, fmgBC, fmgD*, and *fmgE* transcripts in the *csgA* mutant and the induced *csgA* P_*van*_-*fruA* and *csgA* P_*van*_-*fruA D59E* strains during development. Like the *dev* transcript level (Fig. 8B), the *fmg* transcript levels were very low in the *csgA* mutant and in its derivatives induced to produce FruA or FruA D59E (Fig. S18). These findings support the hypothesis that C-signaling activates FruA and that it may do so by a mechanism other than D59 phosphorylation.

To investigate whether LadA regulates *fmg* genes, we measured *fmg* transcript levels in the *ladA* null mutant. Like the *dev* transcript level (Fig. S5A), the *fmgA* and *fmgD* transcript levels were significantly lower in the *ladA* mutant than in WT at 18 h PS (*p* < 0.05 in Student’s two-tailed *t*-tests), but not at 24 or 30 h (Fig. S5B and S5D). The *fmgBC* and *fmgE* transcript levels were significantly lower in the *ladA* mutant at 18 and 24 h, and at all three times, respectively (Fig. S5C and S5E). Hence, LadA does not control the *fmgA* and *fmgD* transcript levels during the period (24 to 30 h PS) when cells commit to spore formation under our conditions, but LadA appears to positively regulate the *fmgBC* transcript level twofold on average at 24 h and the *fmgE* transcript level about fivefold on average at 24 to 30 h.

## Discussion

Our systematic, quantitative analysis of a key circuit in the GRN governing *M. xanthus* fruiting body formation implicates posttranslational regulation of FruA by C-signaling as primarily responsible for *dev* transcript accumulation during the period leading up to and including commitment to spore formation. Mathematical modeling of the *dev* transcript level allowed us to predict the magnitude of potential regulatory mechanisms. Experiments ruled out C-signal-dependent stabilization of *dev* mRNA or highly cooperative binding of FruA and MrpC to two sites in the *dev* promoter region as the explanation for the much higher *dev* transcript level in WT than in the *csgA* mutant. Although the FruA level was twofold lower in the *csgA* mutant than in WT (Fig. 3B and 8A), boosting the FruA or FruA D59E level in the *csgA* mutant had no effect on the *dev* transcript level (Fig. 8B) and did not rescue development (Fig. S15). Taken together, our experimental and computational analyses provide evidence that C-signaling activates FruA at least ninefold posttranslationally during *M. xanthus* development (Fig. 9), and likely by a mechanism other than phosphorylation of D59. The activation of FruA may be considerably greater than ninefold if the *dev* promoter region approaches saturation (Fig. 5C and S12). Since efficient C-signaling requires cells to move into close proximity (Kim & Kaiser, 1990c, Kim & Kaiser, 1990b, Kroos et al., 1988), we propose that activation of FruA by C-signaling acts as a checkpoint for mound formation during the developmental process (Fig. 9).

**Fig. 9.**
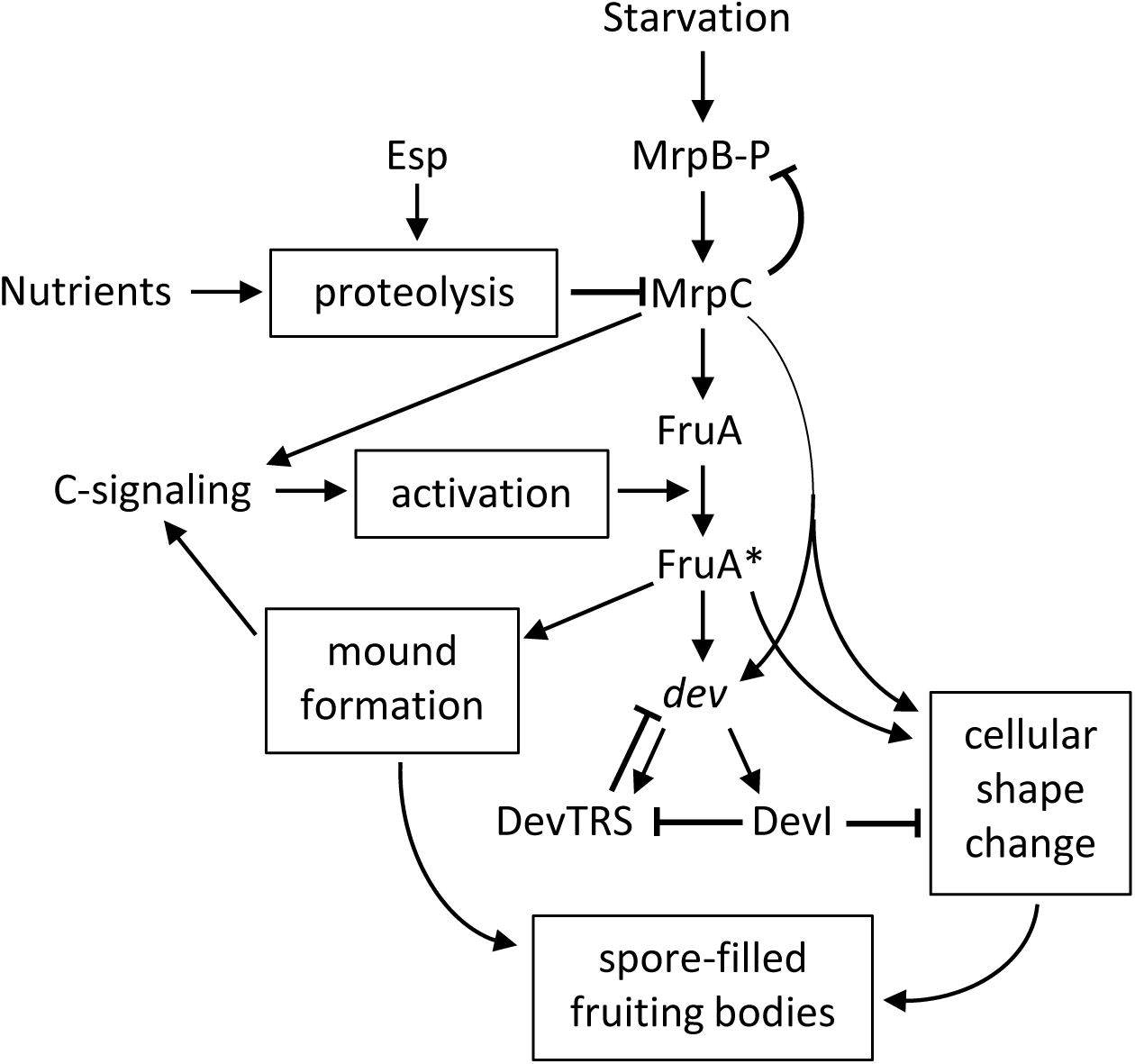
Updated model of the gene regulatory network governing formation of fruiting bodies. Relative to the simplified model shown in Figure 1 (see legend), this model also includes phosphorylated MrpB (MrpB-P) which appears to activate transcription of *mrpC*, and negative autoregulation by MrpC which appears to involve competition with MrpB-P for binding to overlapping sites in the *mrpC* promoter region; proteolysis of MrpC, which is regulated by the Esp signal transduction system that normally slows the developmental process and is regulated by nutrient addition that can halt development; posttranslational activation of FruA to FruA* by C-signaling and promotion of mound formation by FruA*, thus enhancing short-range C-signaling by bringing cells into proximity; the possibility that DevI inhibits negative autoregulation by DevTRS; and speculation that the feed-forward loop involving MrpC and FruA* not only controls transcription of the *dev* operon, but that of genes involved in cellular shape change as well, committing cells to spore formation and resulting in spore-filled fruiting bodies. This model deletes activation of MrpC by C-signaling, which was included as a possibility in Figure 1, but was not supported by our data. See the text for details and references.

### Regulation of FruA by C-signaling

If activation of FruA by C-signaling acts as a checkpoint for mound formation, then active FruA should be present at 18 h PS since mound formation is well underway (Fig. 2). In agreement, mathematical modeling of our data using the assumptions of hypothesis 3 at each time point from 18 to 30 h PS yields a similar result (Fig. S12). This analysis implies that FruA has already been activated by C-signaling at least ninefold by 18 h PS, if the assumptions of hypothesis 3 apply. The assumption that the distal site does not contribute to high cooperativity of MrpC and FruA binding to the *dev* promoter region applies since the *dev* transcript level did not differ significantly in the distal site mutant as compared with WT at 18 or 24 h PS (Fig. 7). We did not measure *dev* transcript stability at 18 to 27 h PS, but at 30 h PS there was no significant difference between WT and the *csgA* mutant (Fig. 6). Therefore, C-signaling may have already activated FruA at least ninefold by 18 h PS, and perhaps as much as 30-fold if the *dev* promoter region approaches saturation (90% saturation in the righthand panel of Fig. S12). We note that during the period from 18 to 30 h PS, the *dev* transcript level rises, but the rise is due to positive autoregulation by DevI (Fig. 4A). Hence, active FruA may not be the limiting factor for *dev* transcription during this period (i.e., the *dev* promoter region may indeed approach saturation binding of active FruA and MrpC). The proximal upstream site in the *dev* promoter region, which is crucial for transcriptional activation, exhibits a higher affinity for cooperative binding of FruA and MrpC than the distal upstream site (Campbell et al., 2015) or several other sites (Robinson et al., 2014, Son et al., 2011), perhaps conferring on *dev* transcription a relatively low threshold for active FruA.

The mechanism of FruA activation by C-signaling is unknown. Since FruA is similar to response regulators of two-component signal transduction systems, phosphorylation by a histidine protein kinase was initially proposed to control FruA activity (Ellehauge et al., 1998, Ogawa et al., 1996). While this potential mechanism of posttranslational control cannot be ruled out, a kinase capable of phosphorylating FruA has not been identified despite considerable effort. Moreover, the atypical receiver domain of FruA and the inability of small-molecule phosphodonors to increase its DNA-binding activity suggest that FruA may not be phosphorylated (Mittal & Kroos, 2009a). In further support of this possibility, we found that expression of FruA D59E, designed to mimic phosphorylation of the receiver domain, did not appear to constitutively activate the protein in the absence of C-signaling (Fig. 8 and S15). Neither did the D59E substitution render FruA unable to be activated, since expression of FruA D59E rescued development of the *fruA* mutant (Fig. S17), consistent a previous study (Ellehauge et al., 1998). Hence, FruA may be an atypical response regulator that is activated by C-signaling *via* a mechanism other than phosphorylation.

Several atypical response regulators have been shown to be active without phosphorylation and a few are regulated by ligand binding (Bourret, 2010, Desai *et al*., 2016). For example, the atypical receiver domain of *Streptomyces venezuelae* JadR1 is bound by jadomycin B, causing JadR1 to dissociate from DNA, and the acylated antibiotic undecylprodigiosin of *Streptomyces coelicolor* may use a similar mechanism to modulate DNA-binding activity of the atypical response regulator RedZ (Wang *et al*., 2009). Conceivably, FruA activity could likewise be regulated by binding of *M. xanthus* diacylglycerols, which have been implicated in C-signaling (Boynton & Shimkets, 2015). Alternatively, FruA could be regulated by a posttranslational modification other than phosphorylation or by binding to another protein (i.e., sequestration).

In addition to regulating FruA activity posttranslationally, C-signaling appears to regulate the FruA level posttranscriptionally. The FruA level was reproducibly twofold lower in the *csgA* mutant than in WT (Fig. 3B and 8A), but the *fruA* transcript level was not significantly different (Fig. 4C and S11A).

These results suggest that positive posttranscriptional regulation of the FruA level requires C-signaling. C-signaling may increase synthesis (i.e., increase *fruA* mRNA accumulation slightly and also increase translation of *fruA* mRNA) and/or decrease turnover of FruA. We did not investigate this further because the FruA deficit in the *csgA* mutant could be overcome with P_*van*_-*fruA* or P_*van*_-*fruA D59E*, yet there was very little effect on the *dev* transcript level (Fig. 8). This demonstrates that the activity of FruA, rather than its level, primarily controls the level of *dev* transcript.

### Regulation by Dev proteins

DevI inhibits sporulation if overexpressed, as in the *devS* mutant (Rajagopalan et al., 2015) (Fig. 2 and S1). Deletion of *devI* or the entire *dev* operon allows spores to begin forming about 6 h earlier than normal, but does not increase the final number of spores (Rajagopalan & Kroos, 2017) (Fig. S1). The level of MrpC was about twofold higher on average in the *devI* mutant than in WT at 15 h PS, perhaps accounting for the observed earlier sporulation, although the difference diminished at 18-24 h PS (Rajagopalan & Kroos, 2017), as reported here (Fig. 3A). It was concluded that DevI may transiently and weakly inhibit translation of *mrpC* transcripts during the period leading up to commitment, delaying sporulation (Rajagopalan & Kroos, 2017). As noted above, DevI positively autoregulates, causing a small rise in the *dev* transcript level by 30 h PS (Fig. 4A, 7, and 8B). Although the mechanism of this feedback loop is unknown, one possibility is that DevI inhibits negative autoregulation by DevTRS (Fig. 9).

In previous studies, mutations in *devT, devR*, or *devS* relieved negative autoregulation, resulting in ∼10-fold higher *dev* transcript accumulation at 24 h PS (Rajagopalan & Kroos, 2017, Rajagopalan et al., 2015). In this study, a *devS* mutant likewise accumulated ∼10-fold more *dev* transcript than WT at 24-30 h PS, and the difference was ∼20-fold at 18 and 21 h PS (Fig. 4A), suggesting that negative autoregulation mediated by DevS has a stronger effect leading up to the commitment period than during commitment. Strong negative autoregulation may promote commitment to sporulation by lowering the level of DevI, which would raise the MrpC level by relieving inhibition of translation of *mrpC* transcripts (Rajagopalan & Kroos, 2017). Our data suggest that negative autoregulation by DevTRS weakens during the commitment period, perhaps accounting for the observed small rise in the *dev* transcript level (Fig. 4A, 7, and 8B). If the elevated *dev* transcript level is accompanied by a small increase in the level of DevI, then DevI may inhibit translation of *mrpC* transcripts, causing the MrpC level to decrease slightly by 30 h PS in WT (Fig. 3A). DevI is predicted to be a 40-residue polypeptide (Rajagopalan et al., 2015) and currently no method has been devised to measure the DevI level. This is a worthwhile goal of future research, as is understanding how cells overcome DevI-mediated inhibition of sporulation (depicted in Fig. 9 as inhibition of cellular shape change).

In addition to regulating the timing of commitment to spore formation, Dev proteins appear to play a role in maturation of spores. Mutations in *dev* genes strongly impair expression of the *exo* operon (Licking *et al*., 2000, Rajagopalan & Kroos, 2017), which encodes proteins that help form the polysaccharide spore coat necessary to maintain cellular shape change and form mature spores (Muller *et al*., 2012, Ueki & Inouye, 2005).

### The role of MrpC

Our results add to a growing list of observations that indicate MrpC functions differently during *M. xanthus* development than originally proposed. We found that MrpC negatively autoregulates accumulation of *mrpC* mRNA about twofold at 18 and 24 h PS (Fig. S8A), and it does so at 18 h PS without significantly altering transcript stability (Fig. S9). This contradicts an earlier study that concluded MrpC positively autoregulates, based on finding that expression of an *mrpC-lacZ* fusion was abolished in an *mrpC* mutant (Sun & Shi, 2001b). Recently, and in agreement with our result, it was reported that MrpC is a negative autoregulator that competes with MrpB for binding to the *mrpC* promoter region (McLaughlin et al., 2018). MrpB, likely when phosphorylated, binds to two sites upstream of the *mrpC* promoter and activates transcription. MrpC binds to multiple sites upstream of the *mrpC* promoter (Nariya & Inouye, 2006, McLaughlin et al., 2018), including two that overlap the MrpB binding sites (McLaughlin et al., 2018). Purified MrpC competes with the MrpB DNA-binding domain for binding to the overlapping sites, supporting a model in which MrpC negatively autoregulates by directly competing with phosphorylated MrpB for binding to overlapping sites (McLaughlin et al., 2018) (Fig. 9).

The role of MrpC in cellular lysis during development appears to be less prominent than originally proposed. MrpC was reported to function as an antitoxin by binding to and inhibiting activity of the MazF toxin protein, an mRNA interferase shown to be important for developmental programmed cell death (Nariya & Inouye, 2008). However, the effect of a null mutation in *mazF* on developmental lysis depends on the presence of a *pilQ1* mutation (Boynton *et al*., 2013, Lee et al., 2012). In *pilQ*+ backgrounds such as our WT strain DK1622, MazF is dispensable for lysis. Here, we found only a slight delay of the *mrpC* mutant relative to WT in terms of the declining number of sonication-sensitive cells at 18-48 h PS (Fig. S11A), comparable to other mutants (*csgA, fruA, devS, csgA* P_*van*_-*fruA, csgA* P_*van*_-*fruA D59E*) that were unable to form spores (Fig. S1 and S15; Table S1). Under our conditions, MrpC appears to play no special role in modulating the cell number during development.

Both the synthesis and the degradation of MrpC are regulated. Synthesis is regulated by phosphorylated MrpB and MrpC acting positively and negatively, respectively, at the level of transcription initiation as described above (McLaughlin et al., 2018) (Fig. 9). Degradation is regulated by the complex Esp signal transduction system (Cho & Zusman, 1999, Higgs *et al*., 2008, Schramm et al., 2012), which presumably senses a signal and controls the activity of an unidentified protease involved in MrpC turnover, thus ensuring that development proceeds at the appropriate pace (Fig. 9). Interestingly, preliminary results suggest that the Esp system does not govern the proteolysis of MrpC observed when nutrients are added at 18 h PS (Rajagopalan & Kroos, 2014) (Y. Hoang, R. Rajagopalan, and L. Kroos; unpublished data). This implies that another system senses nutrients and degrades MrpC to halt development (Fig. 9).

### Combinatorial control by MrpC and FruA

Nutrient-regulated proteolysis of MrpC provides a checkpoint for starvation during the period leading up to and including commitment to sporulation (Rajagopalan & Kroos, 2014) (Fig. 9). If activation of FruA by C-signaling acts as a checkpoint for mound formation as we propose (Fig. 9), then combinatorial control by MrpC and activated FruA could ensure that only starving cells in mounds express genes that commit them to spore formation.

MrpC and FruA bind cooperatively to the promoter regions of five C-signal-dependent genes or operons (*dev, fmgA, fmgBC, fmgD, fmgE*) (Lee et al., 2011, Mittal & Kroos, 2009a, Mittal & Kroos, 2009b, Son et al., 2011, Campbell et al., 2015). In each case, cooperative binding to a site located just upstream of the promoter appears to activate transcription. Hence, MrpC and FruA form a type 1 coherent feed-forward loop with AND logic (Mangan & Alon, 2003). This type of loop is abundant in GRNs and can serve as a sign-sensitive delay element (Mangan & Alon, 2003, Mangan *et al*., 2003). The sign sensitivity refers to a difference in the network response to stimuli in the “OFF to ON” direction versus the “ON to OFF” direction. What this means for the feed-forward loop created by MrpC, FruA, and their target genes is that target gene expression is delayed as MrpC accumulates, awaiting FruA activated by C-signaling (i.e., the “OFF to ON” direction) (Fig. 9). As cells move into mounds and engage in short-range C-signaling, activated FruA would bind cooperatively with MrpC, stimulating transcription of target genes that eventually commit cells to spore formation (depicted in Fig. 9 as cellular shape change). However, if nutrients reappear prior to commitment, MrpC is degraded and transcription of target genes rapidly ceases, halting commitment to sporulation (i.e., the “ON to OFF” direction). The number of target genes may be large since MrpC binds to the promoter regions of hundreds of developmental genes based on ChIP-seq analysis, and in 13 of 15 cases cooperative binding of MrpC and FruA was observed (Robinson et al., 2014).

In addition to the feed-forward loop involving cooperative binding of MrpC and FruA to a site located just upstream of the promoter, the promoter regions of some genes have more complex architectures that confer greater dependence on C-signaling for transcriptional activation. For example, in the *fmgD* promoter region, binding of MrpC to an additional site that overlaps the promoter and the FruA binding site appears to repress transcription, and it has been proposed that a high level of active FruA produced by C-signaling is necessary to outcompete MrpC for binding and result in transcriptional activation (Lee et al., 2011) (Fig. S19A). However, the *fmgD* transcript level in WT did not increase significantly from 18 to 30 h PS (Fig. S5 and S18), suggesting that active FruA is not the limiting factor for transcription during this period. In the *fmgE* promoter region, a distal upstream site with higher affinity for cooperative binding of MrpC and FruA appears to act negatively by competing for binding with the lower affinity site just upstream of the promoter (Son et al., 2011) (Fig. S19B). The *fmgE* transcript level in WT did increase significantly from 18 to 30 h (Fig. S5 and S18), but we cannot conclude that active FruA is the limiting factor since LadA also regulates the *fmgE* transcript level during this period (Fig. S5). In addition to *fmgD* and *fmgE*, other genes depend more strongly on C-signaling and are expressed later during development than *dev* (Kroos & Kaiser, 1987). Some of these genes may require a higher level of active FruA than *dev* and *fmgD* in order to be transcribed. In contrast to the *dev* promoter region, which may have a relatively low threshold for active FruA and therefore approach saturation binding of active FruA and MrpC at 18 h PS (Fig. S12), we predict that the promoter regions of genes essential for commitment to sporulation have more complex architectures and a higher threshold for active FruA. According to this model, C-signal-dependent activation of FruA continues after 18 h PS and the rising level of active FruA triggers commitment beginning at 24 h PS. We speculate that genes governing cellular shape change are under combinatorial control of MrpC and FruA (Fig. 9), and have a high threshold for active FruA.

### Experimental Procedures

#### Bacterial strains, plasmids and primers

The strains, plasmids, and primers used in this study are listed in Table S2. *Escherichia coli* strain DH5α was used for cloning. To construct pSS10, primers FruA-F-NdeI-Gibson and FruA-R-EcoRI-Gibson were used to generate PCR products using chromosomal DNA from *M. xanthus* strain DK1622 as template. The products were combined with NdeI-EcoRI-digested pMR3691 in a Gibson assembly reaction to enzymatically join the overlapping DNA fragments (Gibson *et al*., 2009). The cloned PCR product was verified by DNA sequencing. To construct pSS9, pSS10 was subjected to site-directed mutagenesis using the Quikchange kit (Stratagene) and primers D59E F and D59E R, and the DNA sequence of the *fruA D59E* gene was verified. *M. xanthus* strains with P_*van*_-*fruA* and P_*van*_-*fruA D59E* integrated ectopically were constructed by electroporation (Kashefi & Hartzell, 1995) of pSS10 and pSS9, respectively, selection of transformants on CTT agar containing 15 µg/ml of tetracycline (Iniesta *et al*., 2012), and verification by colony PCR using primers pMR3691 MCS G-F and pMR3691 MCS G-R.

#### *Growth and development of* M. xanthus

Strains of *M. xanthus* were grown at 32°C in CTTYE liquid medium (1% Casitone, 0.2% yeast extract, 10 mMTris-HCl [pH 8.0], 1 mM KH_2_PO_4_-K_2_HPO_4_, 8 mM MgSO_4_ [final pH 7.6]) with shaking at 350 rpm. CTT agar (CTTYE lacking yeast extract and solidified with 1.5% agar) was used for growth on solid medium and was supplemented with 40 µg/ml of kanamycin sulfate or 15 µg/ml of tetracycline as required. Fruiting body development under submerged culture conditions was performed using MC7 (10 mM morpholinepropanesulfonic acid [MOPS; pH 7.0], 1 mM CaCl_2_) as the starvation buffer as described previously (Rajagopalan & Kroos, 2014). Briefly, log-phase CTTYE cultures were centrifuged and cells were resuspended in MC7 at a density of approximately 1,000 Klett units. A 100 μl sample (designated T_0_) was removed, glutaraldehyde (2% final concentration) was added to fix cells, and the sample was stored at 4°C at least 24 h before total cells were quantified as described below. For each developmental sample, 1.5 ml of the cell suspension plus 10.5 ml of MC7 was added to an 8.5-cm-diameter plastic petri plate. Upon incubation at 32°C, cells adhere to the bottom of the plate and undergo development. At the indicated times developing populations were photographed through a microscope and collected as described below.

#### Microscopy

Images of fruiting bodies were obtained using a Leica Wild M8 microscope equipped with an Olympus E-620 digital camera. In order to quantify cells in samples collected and dispersed as described below, high resolution images were obtained with an Olympus BX51 microscope using a differential interference contrast filter and a 40× objective lens, and equipped with an Olympus DP30BW digital camera.

#### Sample collection

At the indicated times the submerged culture supernatant was replaced with 5 ml of fresh MC7 starvation buffer with or without inhibitors as required. Developing cells were scraped from the plate bottom using a sterile cell scraper and the entire contents were collected in a 15-ml centrifuge tube. Samples were mixed thoroughly by repeatedly (three times total) vortexing for 15 s followed by pipetting up and down 15 times. For quantification of total cells, 100 μl of the mixture was removed, glutaraldehyde (2% final concentration) was added to fix cells, and the sample was stored at 4°C for at least 24 h before counting as described below. For measurement of sonication-resistant spores, 400 μl of the mixture was removed and stored at −20°C. For immunoblot analysis, 100 μl of the mixture was added to an equal volume of 2× sample buffer (0.125 M Tris-HCl [pH 6.8], 20% glycerol, 4% sodium dodecyl sulfate [SDS], 0.2% bromophenol blue, 0.2 M dithiothreitol), boiled for 5 min, and stored at −20°C. Immediately after collecting the three samples just described, the remaining 4.4 ml of the developing population was mixed with 0.5 ml of RNase stop solution (5% phenol [pH < 7] in ethanol), followed by rapid cooling in liquid nitrogen until almost frozen, centrifugation at 8,700 × *g* for 10 min at 4°C, removal of the supernatant, freezing of the cell pellet in liquid nitrogen, and storage at −80°C until RNA extraction. Control experiments with a sample collected at 30 h PS indicated that the majority of spores remain intact after boiling in 2× sample buffer or RNA extraction as described below (data not shown), so the proteins and RNAs analyzed are from developing cells that have not yet formed spores.

#### Quantification of total cells and sonication-resistant spores

During development a small percentage of the rod-shaped cells transition to ovoid spores that become sonication-resistant. The number of sonication-resistant spores in developmental samples was quantified as described previously (Rajagopalan & Kroos, 2014). Briefly, each 400-μl sample collected as described above was thawed and sonicated using a model 450 sonifier (Branson) at output setting 2 for 10-s intervals three times with cooling on ice in between. A 60 μl sample was removed and ovoid spores were counted microscopically using 10 μl in a Neubauer counting chamber. A remaining portion of the sample was used to determine total protein concentration as described below. The total cell number, including rod-shaped cells, ovoid spores, and cells in transition between the two, was determined using the glutaraldehyde-fixed samples collected as described above. Each sample was thawed and mixed by vortexing and pipetting, then 10 or 20 μl was diluted with MC7 to 400 μl, sonicated once for 10 s, and all cells were counted microscopically. The total cell number minus the number of sonication-resistant spores was designated the number of sonication-sensitive cells (consisting primarily of rod-shaped cells) and was expressed as a percentage of the total cell number in the corresponding T_0_ sample (consisting only of rod-shaped cells).

#### RNA extraction and analysis

RNA was extracted using the hot-phenol method and the RNA was digested with DNase I (Roche) as described previously (Higgs et al., 2008). One μg of total RNA was subjected to cDNA synthesis using Superscript III reverse transcriptase (InVitrogen) and random primers (Promega), according to the instructions provided by the manufacturers. Control reactions were not subjected to cDNA synthesis. One μl of cDNA at the appropriate dilution (as determined empirically) and 20 pmol of each primer were subjected to qPCR in a 25 μl reaction using 2× reaction buffer (20 mM Tris-HCl [pH 8.3], 13 mM MgCl_2_, 100 mM KCl, 400 μM dNTPs, 4% DMSO, 2× SYBR Green I [Molecular Probes], 0.01% Tween 20, 0.01% NP40, and 0.01 μg/μl of Taq polymerase) as described previously (Bryant *et al*., 2008). qPCR was done in quadruplicate for each cDNA using a LightCycler® 480 System (Roche). A standard curve was generated for each set of qPCRs using *M. xanthus* wild-type strain DK1622 genomic DNA and gene expression was quantified using the relative standard curve method (user bulletin 2; Applied Biosystems). 16S rRNA was used as the internal standard for each sample. Relative transcript levels for mutants are the average of three biological replicates after each replicate was normalized to the transcript level observed for one replicate of WT at 18 h PS in the same experiment. Transcript levels for WT at other times PS were likewise normalized to that observed for WT at 18 h PS in the same experiment. For WT at 18 h PS, the transcript levels of at least three biological replicates from different experiments were normalized to their average, which was set as 1.

#### Immunoblot analysis

A semi-quantitative method of immunoblot analysis was devised to measure the relative levels of MrpC and FruA in many samples collected in different experiments. Equal volumes (10 μl for measurement of MrpC and 15 μl for measurement of FruA) of samples prepared for immunoblot analysis as described above were subjected to SDS-PAGE and immunoblotting as described previously (Rajagopalan & Kroos, 2014, Yoder-Himes & Kroos, 2006). Representative immunoblots for each strain are shown in Figure S20. On each immunoblot, a sample of the wild-type strain DK1622 at 18 h PS served as an internal control for normalization of signal intensities across immunoblots. Signals were detected using a ChemiDoc MP imaging system (Bio-Rad), with exposure times short enough to ensure signals were not saturated, and signal intensities were quantified using Image Lab 5.1 (Bio-Rad) software. After normalization to the internal control, each signal intensity was divided by the total protein concentration of a corresponding sample that had been sonicated for 10-s intervals three times as described above. After removal of a sample for spore quantification, the remaining portion was centrifuged at 10,000 × *g* for 1 min and the total protein concentration of the supernatant was determined using a Bradford (Bradford, 1976) assay kit (Bio-Rad). The resulting values of normalized signal intensity/total protein concentration were further normalized to the average value for all biological replicates of WT at 18 h PS, which was set as 1.

#### Mathematical modeling

##### Activation of dev transcription by FruA and MrpC

FruA and MrpC bind cooperatively to the *dev* promoter region and activate transcription (Campbell et al., 2015). In agreement, no *dev* mRNA was detected in either the *fruA* mutant (Fig. 4A) or the *mrpC* mutant (Fig. 7). We represent the activation of *dev* transcript by FruA and MrpC using a phenomenological Hill’s function,

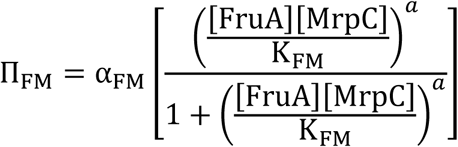

where α_FM_ denotes the maximal *dev* transcription rate, K_FM_ is the half-saturation constant, and a denotes the cooperativity of binding. We model the binding cooperativity using a single Hill exponent for the product of the FruA and MrpC concentrations since one FruA and one MrpC bind cooperatively to the proximal site upstream of the *dev* promoter, based on mobility shift assays (Campbell et al., 2015). Note that this expression will give П_FM_ = 0 when [FruA] = 0 or [MrpC] = 0 (i.e., we have neglected any basal transcription rate as we did not detect *dev* mRNA in the *fruA* or *mrpC* mutant. The expression in brackets can be thought as the promoter occupancy probability (P in the equation below), a dimensional parameter telling what fraction of the promoters will be occupied by the transcription factors for a given value of K_FM_.

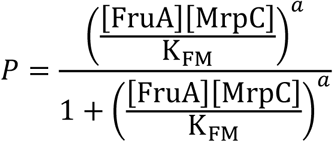

Note that the sensitivity of this expression to changes in the concentrations of FruA and MrpC are maximal when *P*∼0 and minimal near saturation when *P*∼1. In Figure 5 we assess how different hypotheses about the role of C-signaling in *dev* regulation play out at different levels of K_FM_. To facilitate the biological interpretation of the findings, we plot these as a function of *dev* promoter saturation.

##### Feedback regulation by Dev proteins

The *dev* mRNA level is further regulated by Dev proteins DevI and DevS. Our finding that the *dev* transcript level is lower in the *devI* mutant than in WT (Fig. 4A) indicates that DevI is a positive regulator of *dev* mRNA accumulation. In contrast, the *dev* transcript level in the *devS* mutant is significantly higher than in WT (Fig. 4A), indicating that DevS is a negative regulator of *dev* mRNA accumulation. Since the exact mechanisms of regulation by DevI and DevS are unclear, we assume for simplicity that these proteins regulate the *dev* transcript level through independent mechanisms. We model these regulation functions as follows:

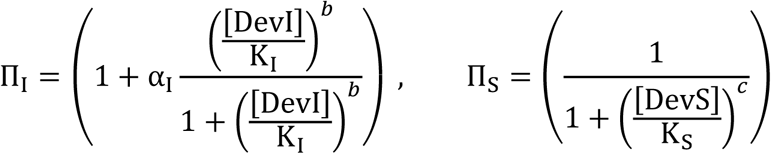

Here, α_I_ is a dimensionless parameter characterizing the feedback strength (i.e., the fold-increase in transcription of the *dev* operon due to DevI), K_I_ is the half-saturation constant, and *b* denotes effective cooperativity (i.e., Hill coefficient) of DevI binding. Likewise, K_S_ is the half-saturation constant and *c* denotes the effective cooperativity of DevS binding. Note that these functions are normalized so that П_I_ = 1 for the *devI* mutant and П_S_ = 1 for the *devS* mutant (i.e., when [DevI] = 0 or [DevS] = 0).

We assume that regulation by the Dev proteins is independent of that by FruA and MrpC, and the effects will be multiplicative:

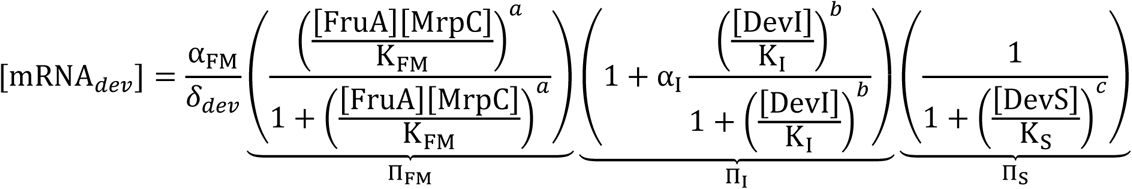

where, K_FM_, K_I_, and K_S_are the saturation constants for regulation by [FruA][MrpC], [DevI], and [DevS], respectively.

##### Numerical procedure to estimate unknown regulation parameters

To explain the difference in the *dev* mRNA level in the *csgA* mutant as compared with WT, in terms of perturbation of potential regulatory mechanisms, we use a mathematical approach where we constrain the FruA ratio ([FruA]_WT_/[FruA]_*cagA*_ ≅ 2) and find the regulation parameters that can result in the observed 22-fold difference in [MRNA_*dev*_]. Specifically, we use the expression of *dev* transcript ratio between WT and the *csgA* mutant below:

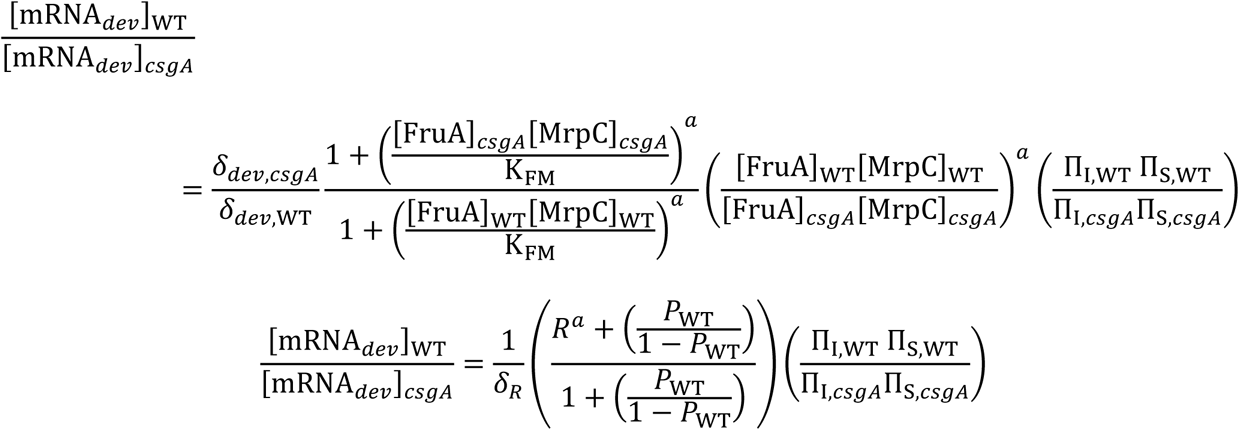

where,

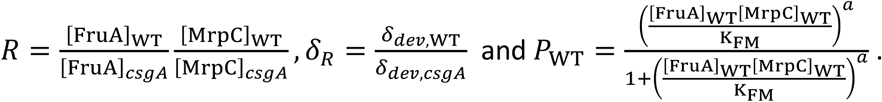

First, we estimate the contribution from Dev protein regulation terms (П_I_, П_S_) in determining the *dev* transcript level in WT and the *csgA* mutant. Since we did not measure the Dev proteins explicitly in our experiments, we estimate their contribution in regulating *dev* transcription in WT by comparing the changes in transcript level in their absence (i.e., in the *devI* and *devS* mutants). Based on our transcript data for WT, and the *devI* and *devS* mutants (Fig. 4A), we have the following relations between the regulation functions; 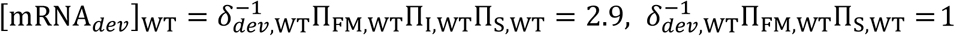 and 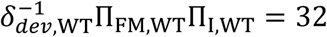. Using these relations, we obtain П_I,WT_ = 2.9, П_S,WT_ = 0.091. For the *csgA* mutant, assuming regulation by Dev proteins is absent due to the low *dev* transcript level, we have П_I,*cagA*_ ≈ 1 and П_S,*cagA*_ ≈ 1. With these estimates, the above expression for *dev* transcript ratio has three unknown parameters *δ*_R_, a, P_WT_.

Next, we determine the required fold change in degradation rate *δ*_R_ for different promoter saturation probability *P*_WT_ values that explains the observed 22-fold difference in *dev* transcript. To estimate this, we set the cooperativity constant (a) to 2 and take the fold change in FruA from the experiments, while assuming MrpC is unchanged between WT and the *csgA* mutant. The result is plotted in Fig. 5A. Then, we determine the required cooperativity a for different *P*_WT_ values with the FruA fold change from the experiments and assuming no change in the degradation rate (*δ*_*R*_ = 1). The result is plotted in Fig. 5B. Finally, we compute the fold change in FruA with *δ*_R_ = 1 and a = 2 for different *P*_WT_ values. The result is shown in Fig. 5C.

### RNA stability

At the indicated time the submerged culture supernatant was replaced with fresh MC7 starvation buffer supplemented with 50 μg/ml of rifampicin to inhibit RNA synthesis. Samples were collected immediately (designated t_0_) and 8 and 16 min later for RNA extraction and analysis as described above, except for each biological replicate the transcript levels after 8 and 16 min were normalized to the transcript level at t_0_, which was set as 1, and the natural log of the resulting values was plotted versus minutes after rifampicin treatment and the slope of a linear fit of the data was used to compute the mRNA half-life.

### *Induction of P*_van_-fruA

To induce expression of *fruA* fused to a vanillate-inducible promoter in *M. xanthus*, the CTTYE growth medium was supplemented with 0.5 mM vanillate when the culture reached 50 Klett units. Growth was continued until the culture reached 100 Klett units, then the culture was centrifuged and cells were resuspended at a density of approximately 1,000 Klett units in MC7 supplemented with 0.5 mM vanillate, followed by submerged culture development as described previously (Rajagopalan & Kroos, 2014).

## Acknowledgements

We thank Monique Floer for advice about high-throughput qPCR and for use of the LightCycler® 480 System. We thank Montserrat Elias-Arnanz and Penelope Higgs for sharing strains. This work was supported by the National Science Foundation (award MCB-1411272) and by salary support for L.K. from Michigan State University AgBioResearch.

## Author contributions

Conception or design of the study: LK, OI, SS, PP

Acquisition of the data: SS, PP

Analysis or interpretation of the data: SS, PP, LK, OI

Writing of the manuscript: LK, SS, PP, OI

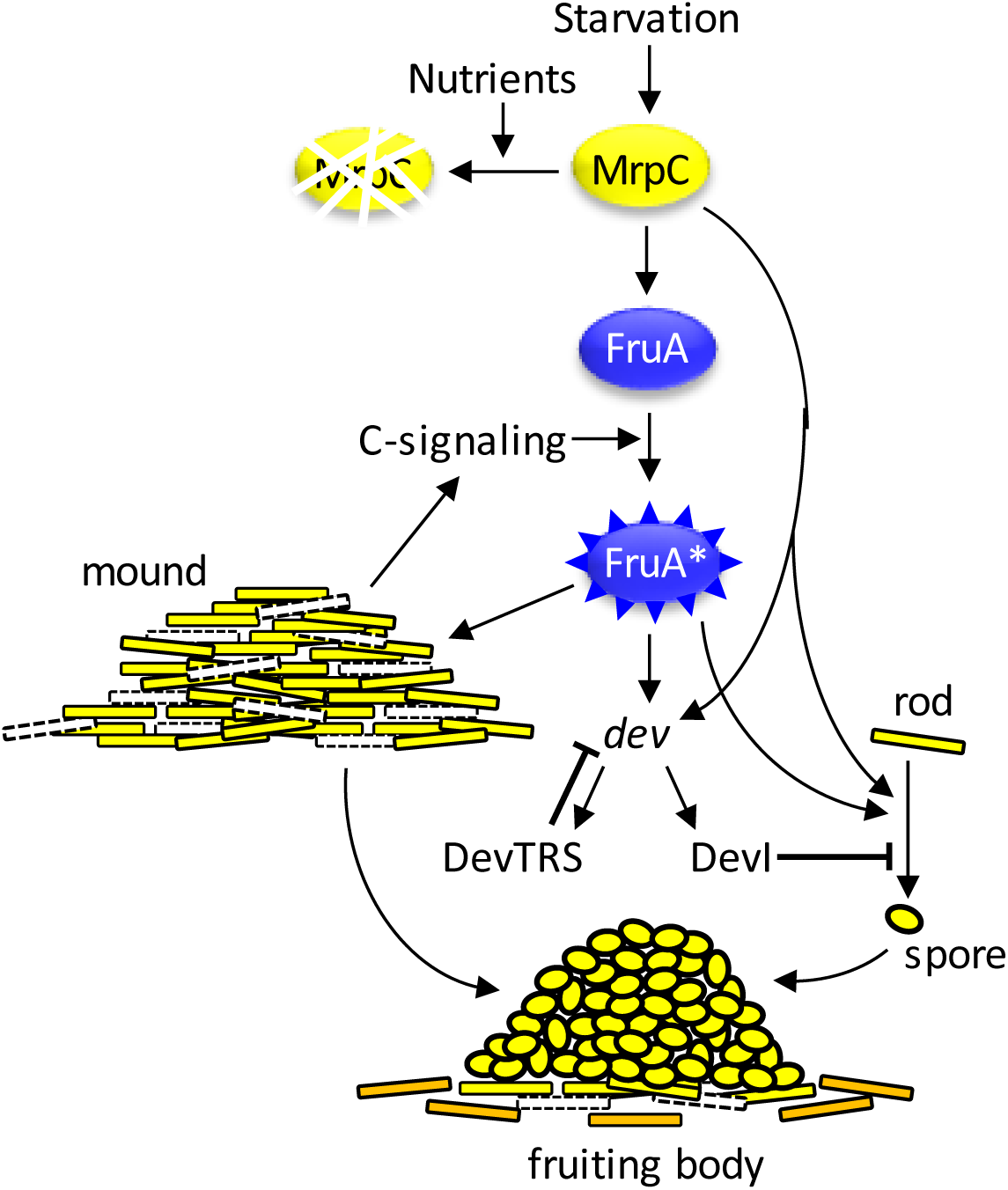
Graphical abstract.

## Abbreviated summary

Starvation promotes MrpC accumulation, whereas nutrients favor proteolysis. MrpC activates transcription of *fruA*, but FruA protein appears to be activated by short-range C-signaling in a cycle leading to mound formation and lysis of some cells. Activated FruA* and MrpC are proposed to cooperatively stimulate transcription of the *dev* operon and genes that commit starving rod-shaped cells to form spores, while Dev proteins slow commitment, resulting in a spore-filled fruiting body surrounded by peripheral rods.

## References

Barratt, M. J., C. Lebrilla, H. Y. Shapiro & J. I. Gordon, (2017) The gut microbiota, food science, and human nutrition: a timely marriage. Cell Host Microbe 22: 134–141.

Bourret, R. B., (2010) Receiver domain structure and function in response regulator proteins. Curr. Opin. Microbiol. 13: 142–149.

Boynton, T. O., J. L. McMurry & L. J. Shimkets, (2013) Characterization of *Myxococcus xanthus* MazF and implications for a new point of regulation. Mol. Microbiol. 87: 1267–1276.

Boynton, T. O. & L. J. Shimkets, (2015) *Myxococcus* CsgA, *Drosophila* Sniffer and human HSD17B10 are cardiolipin phospholipases. Genes Dev. 29: 1903–1914.

Boysen, A., E. Ellehauge, B. Julien & L. Sogaard-Andersen, (2002) The DevT protein stimulates synthesis of FruA, a signal transduction protein required for fruiting body morphogenesis in *Myxococcus xanthus*. J. Bacteriol. 184: 1540–1546.

Bradford, M., (1976) A rapid and sensitive method for the quantitation of microgram quantities of protein utilizing the principle of protein-dye binding. Anal. Biochem. 72: 248–254.

Bretl, D. J. & J. R. Kirby, (2016) Molecular mechanisms of signaling in *Myxococcus xanthus* development. J. Mol. Biol. 428: 3805–3830.

Bryant, G. O., V. Prabhu, M. Floer, X. Wang, D. Spagna, D. Schreiber & M. Ptashne, (2008) Activator control of nucleosome occupancy in activation and repression of transcription. PLoS Biol. 6: 2928–2939.

Bush, M. J., N. Tschowri, S. Schlimpert, K. Flardh & M. J. Buttner, (2015) c-di-GMP signalling and the regulation of developmental transitions in streptomycetes. Nat. Rev. Microbiol. 13: 749–760.

Campbell, A., P. Viswanathan, T. Barrett, B. Son, S. Saha & L. Kroos, (2015) Combinatorial regulation of the *dev* operon by MrpC2 and FruA during *Myxococcus xanthus* development. J. Bacteriol. 197: 240–251.

Cho, K. & D. R. Zusman, (1999) Sporulation timing in *Myxococcus xanthus* is contolled by the *espAB* locus. Mol. Microbiol. 34: 714–725.

Davidson, E. H. & M. S. Levine, (2008) Properties of developmental gene regulatory networks. Proc. Natl. Acad. Sci. USA 105: 20063–20066.

Davlieva, M., Y. Shi, P. G. Leonard, T. A. Johnson, M. R. Zianni, C. A. Arias, J. E. Ladbury & Y. Shamoo, (2015) A variable DNA recognition site organization establishes the LiaR-mediated cell envelope stress response of enterococci to daptomycin. Nucleic Acids Res. 43: 4758–4773.

Desai, S. K., R. S. Winardhi, S. Periasamy, M. M. Dykas, Y. Jie & L. J. Kenney, (2016) The horizontally-acquired response regulator SsrB drives a *Salmonella* lifestyle switch by relieving biofilm silencing. eLife 5: e10747.

Domian, I. J., K. C. Quon & L. Shapiro, (1997) Cell type-specific phosphorylation and proteolysis of a transcriptional regulator controls the G1-to-S transition in a bacterial cell cycle. Cell 90: 415–424.

Drapek, C., E. E. Sparks & P. N. Benfey, (2017) Uncovering gene regulatory networks controlling plant cell differentiation. Trends Genet. 33: 529–539.

Ellehauge, E., M. Norregaard-Madsen & L. Sogaard-Andersen, (1998) The FruA signal transduction protein provides a checkpoint for the temporal co-ordination of intercellular signals in *Myxococcus xanthus* development. Mol. Microbiol. 30: 807–817.

Frum, T. & A. Ralston, (2015) Cell signaling and transcription factors regulating cell fate during formation of the mouse blastocyst. Trends Genet. 31: 402–410.

Galperin, M. Y., (2010) Diversity of structure and function of response regulator output domains. Curr. Opin. Microbiol. 13: 150–159.

Gibson, D. G., L. Young, R. Y. Chuang, J. C. Venter, C. A. Hutchison, 3rd & H. O. Smith, (2009) Enzymatic assembly of DNA molecules up to several hundred kilobases. Nat. Methods 6: 343–345.

Hagen, T. J. & L. J. Shimkets, (1990) Nucleotide sequence and transcriptional products of the *csg* locus of *Myxococcus xanthus*. J. Bacteriol. 172: 15–23.

Higgs, P. I., S. Jagadeesan, P. Mann & D. R. Zusman, (2008) EspA, an orphan hybrid histidine protein kinase, regulates the timing of expression of key developmental proteins of *Myxococcus xanthus*. J. Bacteriol. 190: 4416–4426.

Iniesta, A. A., F. Garcia-Heras, J. Abellon-Ruiz, A. Gallego-Garcia & M. Elias-Arnanz, (2012) Two systems for conditional gene expression in *Myxococcus xanthus* inducible by isopropyl-?-D-thiogalactopyranoside or vanillate. J. Bacteriol. 194: 5875–5885.

Jansson, J. K. & K. S. Hofmockel, (2018) The soil microbiome-from metagenomics to metaphenomics. Curr. Opin. Microbiol. 43: 162–168.

Kashefi, K. & P. Hartzell, (1995) Genetic supression and phenotypic masking of a *Myxococcus xanthux frzF*defect. Molec. Microbiol. 15: 483–494.

Kim, S. K. & D. Kaiser, (1990a) C-factor: a cell-cell signaling protein required for fruiting body morphogenesis of *M. xanthus*. Cell 61: 19–26.

Kim, S. K. & D. Kaiser, (1990b) Cell alignment required in differentiation of *Myxococcus xanthus*. Science 249: 926–928.

Kim, S. K. & D. Kaiser, (1990c) Cell motility is required for the transmission of C-factor, an intercellular signal that coordinates fruiting body morphogenesis of *Myxococcus xanthus*. Genes Dev. 4: 896–905.

Klose, K. E., D. S. Weiss & S. Kustu, (1993) Glutamate at the site of phosphorylation of nitrogen-regulatory protein NTRC mimics aspartyl-phosphate and activates the protein. J. Mol. Biol. 232: 67–78.

Kroos, L., (2017) Highly signal-responsive gene regulatory network governing *Myxococcus* development. Trends Genet. 33: 3–15.

Kroos, L., P. Hartzell, K. Stephens & D. Kaiser, (1988) A link between cell movement and gene expression argues that motility is required for cell-cell signaling during fruiting body development. Genes Dev. 2: 1677–1685.

Kroos, L. & D. Kaiser, (1987) Expression of many developmentally regulated genes in *Myxococcus* depends on a sequence of cell interactions. Genes Dev. 1: 840–854.

Lan, C. Y. & M. M. Igo, (1998) Differential expression of the OmpF and OmpC porin proteins in *Escherichia coli* K-12 depends upon the level of active OmpR. J. Bacteriol. 180: 171–174.

Lee, B., C. Holkenbrink, A. Treuner-Lange & P. I. Higgs, (2012) *Myxococcus xanthus* developmental cell fate production: heterogeneous accumulation of developmental regulatory proteins and reexamination of the role of MazF in developmental lysis. J. Bacteriol. 194: 3058–3068.

Lee, J., B. Son, P. Viswanathan, P. Luethy & L. Kroos, (2011) Combinatorial regulation of *fmgD* by MrpC2 and FruA during *Myxococcus xanthus* development. J. Bacteriol. 193: 1681–1689.

Licking, E., L. Gorski & D. Kaiser, (2000) A common step for changing cell shape in fruiting body and starvation-independent sporulation of *Myxococcus xanthus*. J. Bacteriol. 182: 3553–3558.

Lobedanz, S. & L. Sogaard-Andersen, (2003) Identification of the C-signal, a contact-dependent morphogen coordinating multiple developmental responses in *Myxococcus xanthus*. Genes Dev. 17: 2151–2161.

Mangan, S. & U. Alon, (2003) Structure and function of the feed-forward loop network motif. Proc. Natl. Acad. Sci. USA 100: 11980–11985.

Mangan, S., A. Zaslaver & U. Alon, (2003) The coherent feedforward loop serves as a sign-sensitive delay element in transcription networks. J. Mol. Biol. 334: 197–204.

McLaughlin, P. T., V. Bhardwaj, B. E. Feeley & P. I. Higgs, (2018) MrpC, a CRP/Fnr homolog, functions as a negative autoregulator during the *Myxococcus xanthus* multicellular developmental program. Mol. Microbiol. 109: 245–261.

Mittal, S. & L. Kroos, (2009a) A combination of unusual transcription factors binds cooperatively to control *Myxococcus xanthus* developmental gene expression. Proc. Natl. Acad. Sci. USA 106: 1965–1970.

Mittal, S. & L. Kroos, (2009b) Combinatorial regulation by a novel arrangement of FruA and MrpC2 transcription factors during *Myxococcus xanthus* development. J. Bacteriol. 191: 2753–2763.

Muller, F. D., C. W. Schink, E. Hoiczyk, E. Cserti & P. I. Higgs, (2012) Spore formation in *Myxococcus xanthus* is tied to cytoskeleton functions and polysaccharide spore coat deposition. Mol. Microbiol. 83: 486–505.

Nariya, H. & M. Inouye, (2008) MazF, an mRNA interferase, mediates programmed cell death during multicellular *Myxococcus* development. Cell 132: 55–66.

Nariya, H. & S. Inouye, (2005) Identification of a protein Ser/Thr kinase cascade that regulates essential transcriptional activators in *Myxococcus xanthus* development. Mol. Microbiol. 58: 367–379.

Nariya, H. & S. Inouye, (2006) A protein Ser/Thr kinase cascade negatively regulates the DNA-binding activity of MrpC, a smaller form of which may be necessary for the *Myxococcus xanthus* development. Mol. Microbiol. 60: 1205–1217.

Norman, T. M., N. D. Lord, J. Paulsson & R. Losick, (2015) Stochastic switching of cell fate in microbes. Annu. Rev. Microbiol. 69: 381–403.

O’Connor, K. A. & D. R. Zusman, (1991) Development in *Myxococcus xanthus* involves differentiation into two cell types, peripheral rods and spores. J. Bacteriol. 173: 3318–3333.

Ogawa, M., S. Fujitani, X. Mao, S. Inouye & T. Komano, (1996) FruA, a putative transcription factor essential for the development of *Myxococcus xanthus*. Mol. Microbiol. 22: 757–767.

Rajagopalan, R. & L. Kroos, (2014) Nutrient-regulated proteolysis of MrpC halts expression of genes important for commitment to sporulation during *Myxococcus xanthus* development. J. Bacteriol. 196: 2736–2747.

Rajagopalan, R. & L. Kroos, (2017) The *dev* operon regulates the timing of sporulation during *Myxococcus xanthus* development. J. Bacteriol. 199: e00788–00716.

Rajagopalan, R., S. Wielgoss, G. Lippert, G. J. Velicer & L. Kroos, (2015) *devI* is an evolutionarily young negative regulator of *Myxococcus xanthus* development. J Bacteriol 197: 1249–1262.

Robinson, M., B. Son & L. Kroos, (2014) Transcription factor MrpC binds to promoter regions of many developmentally-regulated genes in *Myxococcus xanthus*. BMC Genomics 15: 1123.

Rolbetzki, A., M. Ammon, V. Jakovljevic, A. Konovalova & L. Sogaard-Andersen, (2008) Regulated secretion of a protease activates intercellular signaling during fruiting body formation in *M. xanthus*. Dev. Cell 15: 627–634.

Sager, B. & D. Kaiser, (1993) Two cell-density domains within the *Myxococcus xanthus* fruiting body. Proc. Natl. Acad. Sci. USA 90: 3690–3694.

Schramm, A., B. Lee & P. I. Higgs, (2012) Intra-and inter-protein phosphorylation between two hybrid histidine kinases controls *Myxococcus xanthus* developmental progression. J. Biol. Chem. 287: 25060–25072.

Shimkets, L. J., R. E. Gill & D. Kaiser, (1983) Developmental cell interactions in *Myxococcus xanthus* and the *spoC* locus. Proc. Natl. Acad. Sci. USA 80: 1406–1410.

Son, B., Y. Liu & L. Kroos, (2011) Combinatorial regulation by MrpC2 and FruA involves three sites in the *fmgE* promoter region during *Myxococcus xanthus* development. J. Bacteriol. 193: 2756–2766.

Srinivasan, D. & L. Kroos, (2004) Mutational analysis of the *fruA* promoter region demonstrates that C-box and 5-base-pair elements are important for expression of an essential developmental gene of *Myxococcus xanthus*. J. Bacteriol. 186: 5961–5967.

Stock, A. M., V. L. Robinson & P. N. Goudreau, (2000) Two-component signal transduction. Annu. Rev. Biochem. 69: 183–215.

Sun, H. & W. Shi, (2001a) Analyses of *mrp* genes during *Myxococcus xanthus* development. J. Bacteriol. 183: 6733–6739.

Sun, H. & W. Shi, (2001b) Genetic studies of *mrp*, a locus essential for cellular aggregation and sporulation of *Myxococcus xanthus*. J. Bacteriol. 183: 4786–4795.

Thony-Meyer, L. & D. Kaiser, (1993) *devRS*, an autoregulated and essential genetic locus for fruiting body development in *Myxococcus xanthus*. J. Bacteriol. 175: 7450–7462.

Ueki, T. & S. Inouye, (2003) Identification of an activator protein required for the induction of *fruA*, a gene essential for fruiting body development in *Myxococcus xanthus*. Proc. Natl. Acad. Sci. USA 100: 8782–8787.

Ueki, T. & S. Inouye, (2005) Identification of a gene involved in polysaccharide export as a transcription target of FruA, an essential factor for *Myxococcus xanthus* development. J. Biol. Chem. 280: 32279–32284.

van Gestel, J., H. Vlamakis & R. Kolter, (2015) Division of labor in biofilms: the ecology of cell differentiation. Microbiol. Spectr. 3: MB–0002-2014.

Viswanathan, P., K. Murphy, B. Julien, A. G. Garza & L. Kroos, (2007a) Regulation of *dev*, an operon that includes genes essential for *Myxococcus xanthus* development and CRISPR-associated genes and repeats. J. Bacteriol. 189: 3738–3750.

Viswanathan, P., T. Ueki, S. Inouye & L. Kroos, (2007b) Combinatorial regulation of genes essential for *Myxococcus xanthus* development involves a response regulator and a LysR-type regulator. Proc. Natl. Acad. Sci. USA 104: 7969–7974.

Wang, L., X. Tian, J. Wang, H. Yang, K. Fan, G. Xu, K. Yang & H. Tan, (2009) Autoregulation of antibiotic biosynthesis by binding of the end product to an atypical response regulator. Proc Natl Acad Sci U S A 106: 8617–8622.

Yang, Z. & P. Higgs, (2014) Myxobacteria: genomics, cellular and molecular biology. In. Norfolk, UK: Caister Academic Press, pp.

Yoder-Himes, D. & L. Kroos, (2006) Regulation of the *Myxococcus xanthus* C-signal-dependent ?4400 promoter by the essential developmental protein FruA. J. Bacteriol. 188: 5167–5176.

