## Supplemental files for "Systematic analysis of the *Myxococcus xanthus* developmental gene regulatory network supports posttranslational regulation of FruA by C-signaling"

**Table S1** Cell and spore numbers

| Strain | Sonication-sensitive cells at $T_0$ ($10^7$/mL) | Sonication-resistant spores at $T_{48}$ ($10^7$/mL) |
|---|---|---|
| wild type | 140 ± 16 | 1.5 ± 0.3 |
| *csgA* | 150 ± 30 | < 0.05 |
| *fruA* | 150 ± 25 | < 0.05 |
| *devI* | 150 ± 27 | 2 ± 0.4 |
| *devS* | 140 ± 17 | < 0.05 |
| *ladA* | 150 ± 5 | 0.7 ± 0.3 |
| distal site | 150 ± 6 | 1.4 ± 0.2 |
| *mrpC* | 130 ± 16 | < 0.05 |
| *csgA* $P_{van}$-*fruA* | 150 ± 7 | < 0.05 |
| *csgA* $P_{van}$-*fruA D59E* | 140 ± 8 | < 0.05 |

Wild-type DK1622 and its indicated mutant derivatives were subjected to starvation under submerged culture conditions. The number of rod-shaped sonication-sensitive cells at $T_0$ and the number of sonication-resistant spores at 48 h poststarvation were counted microscopically using a Neubauer chamber. Values indicate the average of at least 3 biological replicates and one standard deviation.

**Table S2** Strains, plasmids, and primers

| **Bacterial strain, plasmid, or primer** | **Description** | **Source or reference** |
|---|---|---|
| **Strains** | | |
| *E. coli* | | |
| DH5α | $\lambda^-$ $\phi$80dlacZΔM15 Δ(*lacZYA-argF*)*U169 recA1 endA1 hsdR17*($r_K^-$ $m_K^-$) *supE44 thi-1 gyrA relA1* | (Hanahan, 1983) |
| *M. xanthus* | | |
| DK1622 | Laboratory strain | (Kaiser, 1979) |
| SW2808 | Δ*mrpC* | (Sun & Shi, 2001) |
| DK5285 | *fruA*::Tn*5 lac* Ω4491 ($Km^r$) | (Kroos & Kaiser, 1987) |
| DK11209 | Δ*devS* | (Viswanathan *et al.*, 2007a) |
| MRR7 | Δ*devI* | (Rajagopalan *et al.*, 2015) |
| DK5208 | *csgA*::Tn*5-132* Ω205 ($Tc^r$) | (Shimkets & Asher, 1988) |
| MRR33 | *csgA*::pRR028 ($Km^r$) | (Rajagopalan & Kroos, 2017) |
| MPVlysR | *ladA*::pPV4330KO | (Viswanathan *et al.*, 2007b) |
| MSS1 | A deletion of chromosomal DNA between positions -254 and -228 relative to the *dev* transcriptional start site | (Campbell *et al.*, 2015) |
| MSS3 | *csgA*::pRR028 ($Km^r$) MXAN_0018-MXAN_0019::pSS10 ($Tc^r$) | This study |
| MSS5 | *csgA*::pRR028 ($Km^r$) MXAN_0018-MXAN_0019::pSS9 ($Tc^r$) | This study |
| MSS6 | *fruA*::Tn*5 lac* Ω4491 ($Km^r$) MXAN_0018-MXAN_0019::pSS10 ($Tc^r$) | This study |
| MSS7 | *fruA*::Tn*5 lac* Ω4491 ($Km^r$) MXAN_0018-MXAN_0019::pSS9 ($Tc^r$) | This study |
| **Plasmids** | | |
| pSS10 | $Tc^r$; pMR3691 with *fruA* inserted at MCS_G | This study |
| pSS9 | $Tc^r$; pMR3691 with *fruA D59E* inserted at MCS_G | This study |
| pMR3691 | $Tc^r$; *M. xanthus* MXAN_0018-MXAN_0019-$P_{R3-4}$::*vanR*-$P_{van}$-MCS_G | (Iniesta *et al.*, 2012) |
| **Primers** | | |
| FruA-F-NdeI-Gibson | GATGCGAGGAAACGCATATGGCAACCAATCAAGCAGCGATTCGTG | This study |
| FruA-R-EcoRI-Gibson | GTACGCGTAACGTTCGAATTCCTAGAGGTCCGGCGGCGGCCGGA | This study |
| pMR3691 MCS G-F | CACGATGCGAGGAAACGCA | This study |
| pMR3691 MCS G-R | CACCGGTACGCGTAACGTTC | This study |
| 16S rRNA fwd | CAAGGGAACTGAGAGACAGG | (Ossa *et al.*, 2007) |

| | | |
|---|---|---|
| 16S rRNA rev | CTCTAGAGATCCACTACTTGCG | (Ossa et al., 2007) |
| fruA oPH252 | CGTCACGGAAGGCATCAATC | (Rajagopalan & Kroos, 2017) |
| fruA oPH253 | CGAGATGATTTCCGGTGTGC | (Rajagopalan & Kroos, 2017) |
| mrpC qPCR F | GGAGGCCATCGACTTCAAGG | (Rajagopalan & Kroos, 2014) |
| mrpC qPCR R | GGCCGGACTTCAGCAGGTAG | (Rajagopalan & Kroos, 2014) |
| cas6-F | TGGGGAAATCTAATGGTGTTTG | This study |
| cas6-R | GAGAACAGCAGATAGGCATGGT | This study |
| D59E F | CCGCAGGTCGCGGTGATGGAGGTGGAGGGCGACAG CGAG | This study |
| D59E R | CTCGCTGTCGCCCTCCACCTCCATCACCGCGACCTGC GG | This study |
| FmgA-F9 | AAGACGCGCATCAAGGACG | This study |
| FmgA-R9 | CCAGACTTCGAAGCCATCCGAG | This study |
| FmgB-F3 | TGCGCTGCTGTACGACTCC | This study |
| FmgB-R3 | GATGGCCTGGACGGGGCA | This study |
| FmgD-F3N | TTACGGTGGCACCGCATTC | This study |
| FmgD-R3N | CTGGGCTTCCGTCATCTTG | This study |
| FmgE-F3N | CTCATCTGTCGCGGCCAA | This study |
| FmgE-R3N | ACAGCGGTCAGTTCTGAATG | This study |

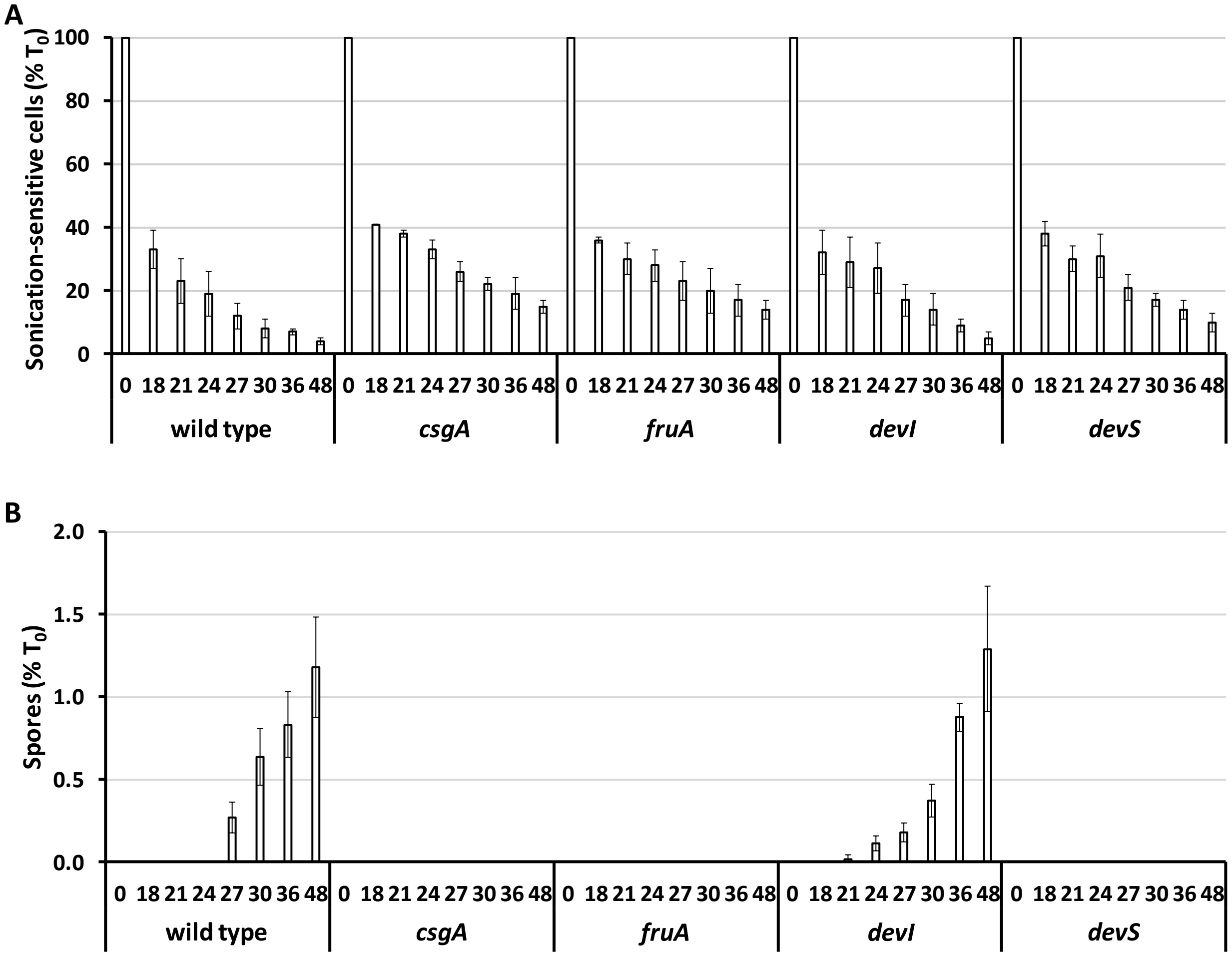


Figure S1. Cellular changes during *M. xanthus* development. Wild-type DK1622 and its indicated mutant derivatives were subjected to starvation under submerged culture conditions and samples were collected at the indicated number of hours poststarvation for quantification of (A) sonication-sensitive cells and (B) sonication-resistant spores. Values are expressed as a percentage of the number of rod-shaped cells present at the time starvation initiated development ($T_0$) (Table S1). Bars show the average of three biological replicates and error bars show one standard deviation.

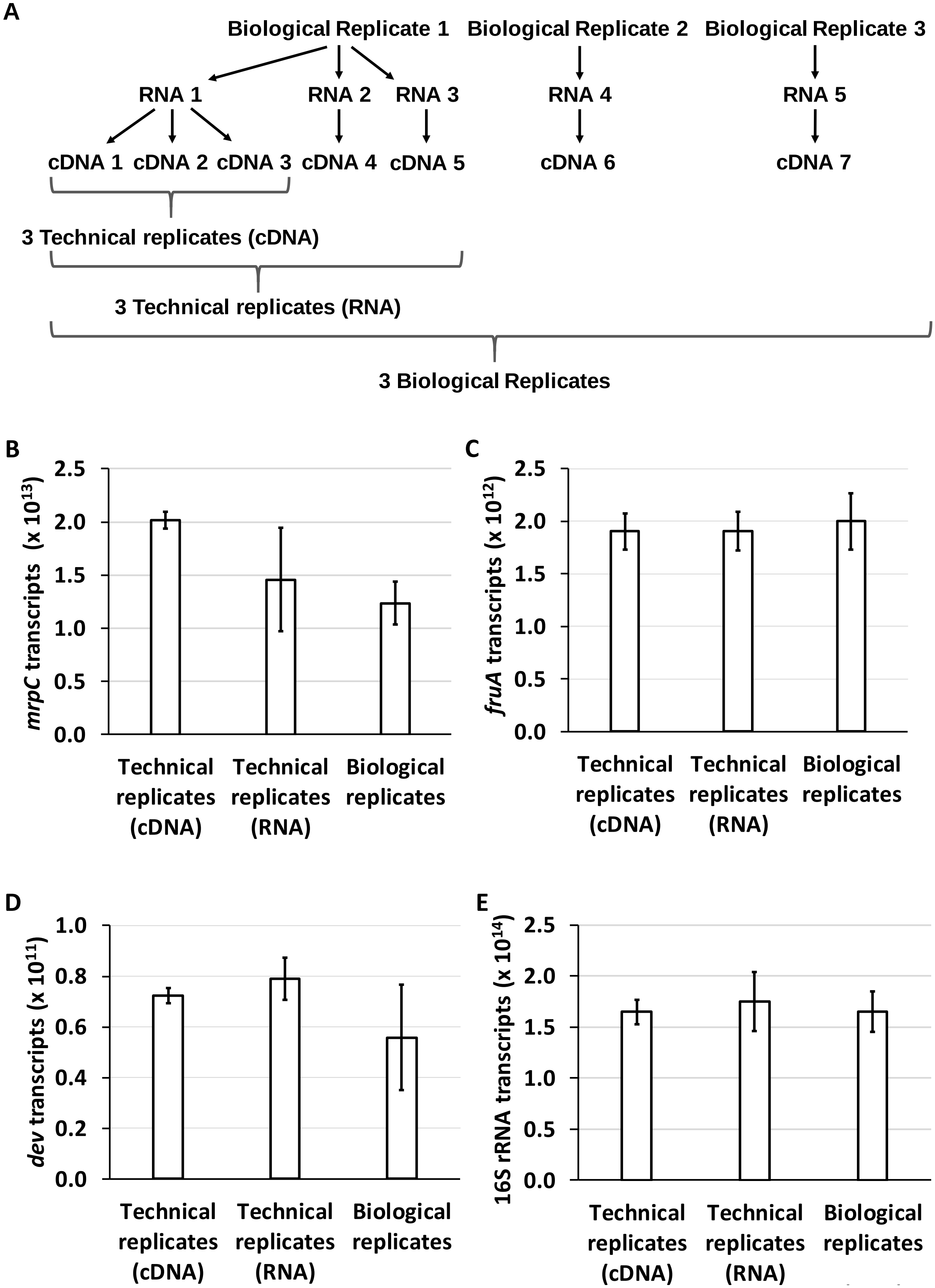
A
Biological Replicate 1
Biological Replicate 2
Biological Replicate 3
RNA 1
RNA 2
RNA 3
RNA 4
RNA 5
cDNA 1
cDNA 2
cDNA 3
cDNA 4
cDNA 5
cDNA 6
cDNA 7
3 Technical replicates (cDNA)
3 Technical replicates (RNA)
3 Biological Replicates
B
mrpC transcripts (x 10^13)
2.5
2.0
1.5
1.0
0.5
0.0
Technical replicates (cDNA)
Technical replicates (RNA)
Biological replicates
C
fruA transcripts (x 10^12)
2.5
2.0
1.5
1.0
0.5
0.0
Technical replicates (cDNA)
Technical replicates (RNA)
Biological replicates
D
dev transcripts (x 10^11)
1.0
0.8
0.6
0.4
0.2
0.0
Technical replicates (cDNA)
Technical replicates (RNA)
Biological replicates
E
16S rRNA transcripts (x 10^14)
2.5
2.0
1.5
1.0
0.5
0.0
Technical replicates (cDNA)
Technical replicates (RNA)
Biological replicates

Figure S2. Reproducibility of RNA measurements. (A) Experimental scheme. Three biological replicates of wild-type DK1622 were subjected to starvation under submerged culture conditions and samples were collected at 24 h poststarvation. One biological replicate sample was used to prepare RNA in triplicate and one of these RNA samples was used to prepare cDNA in triplicate. (B-E) Variation in transcript numbers among cDNA technical replicates, RNA technical replicates (the average of cDNA technical replicates was used as one of the values), and biological replicates. Transcript numbers are per μg total RNA. Bars show the average and error bars show one standard deviation.

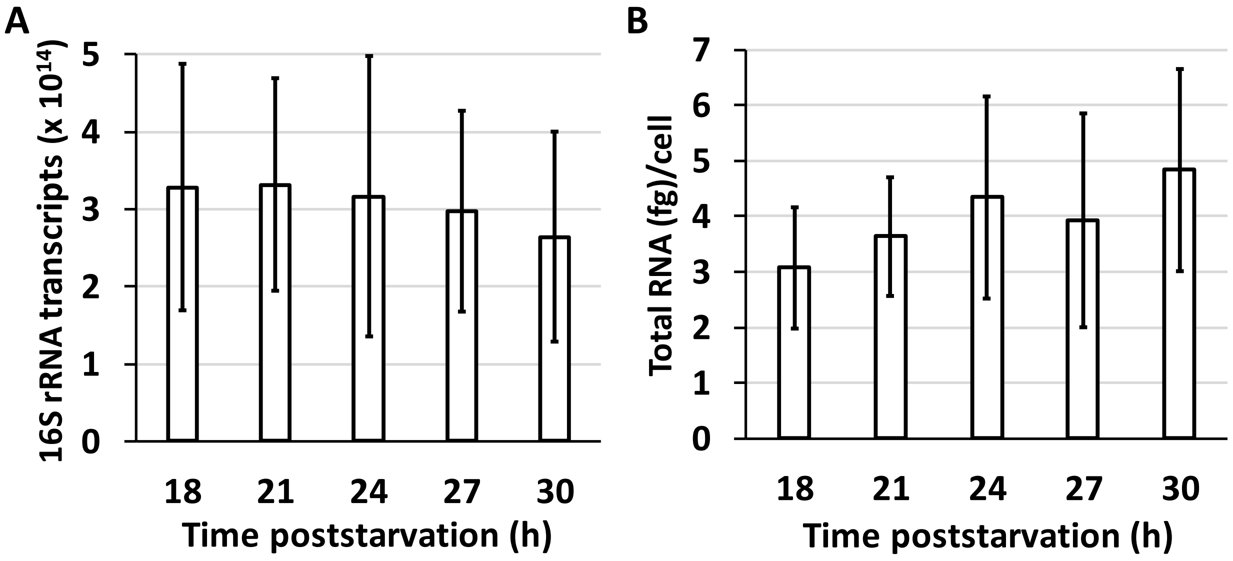


Figure S3. Validation of 16S rRNA as an internal standard for RT-qPCR analysis during *M. xanthus* development. Four biological replicates of wild-type DK1622 were subjected to starvation under submerged culture conditions and RNA was prepared from samples collected at the indicated times poststarvation. (A) Transcript numbers per μg total RNA. (B) Total RNA yield per cell. The RNA yield in femtograms (fg) was divided by the number of rod-shaped cells in the sample prior to RNA preparation. Bars show the average and error bars show one standard deviation.

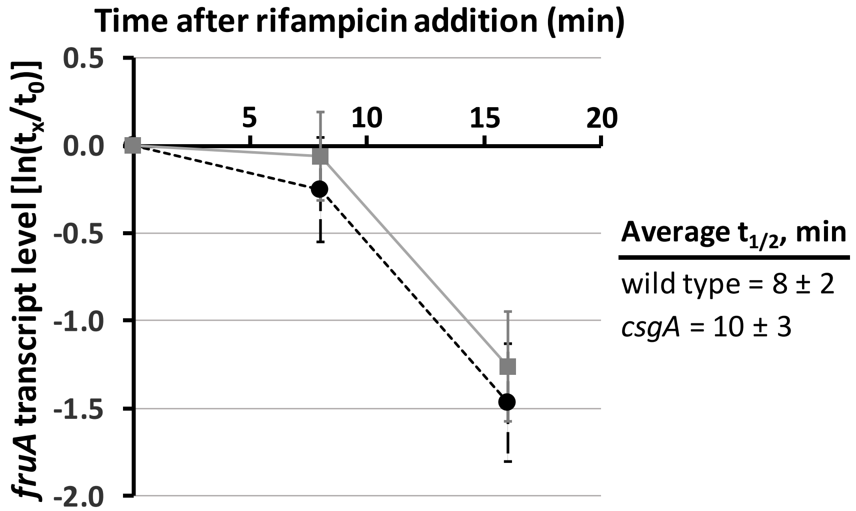


Figure S4. *fruA* transcript stability. Wild-type DK1622 and the *csgA* mutant were subjected to starvation under submerged culture conditions for 30 h. The overlay was replaced with fresh starvation buffer containing rifampicin (50 μg/ml) and samples were collected immediately ($t_0$) and at the times indicated ($t_x$) for measurement of the *fruA* transcript level by RT-qPCR. Transcript levels at $t_x$ were normalized to that at $t_0$ for each of three biological replicates and used to determine the transcript half-life for each replicate. The average half-life (Average $t_{1/2}$) and one standard deviation are shown, and the difference is not statistically significant ($p = 0.42$ in a Student's two-tailed *t*-test). The graph shows the average $\ln(t_x/ t_0)$ and one standard deviation for the three biological replicates of wild type (black dashed line) and the *csgA* mutant (gray solid line).

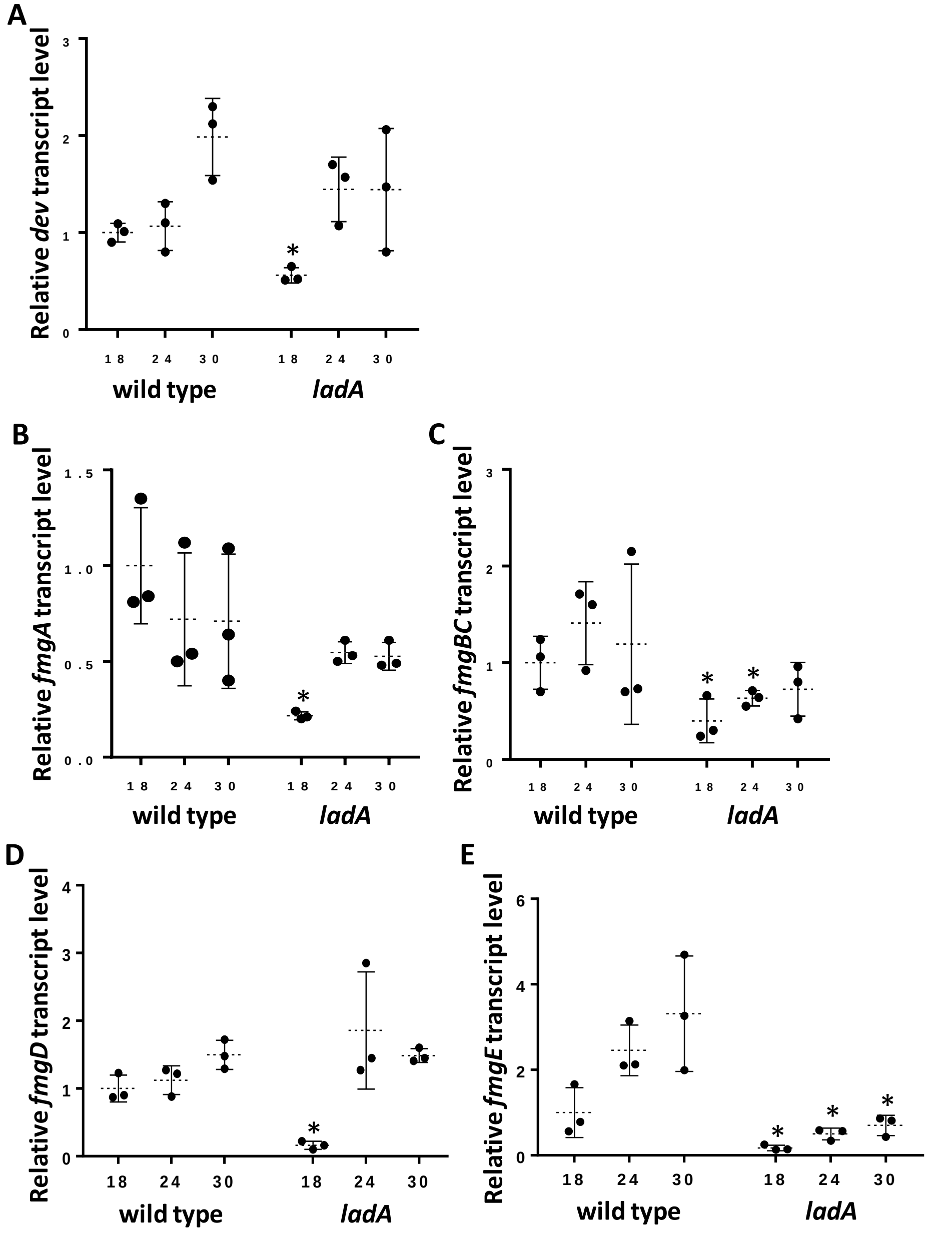


Figure S5. Levels of *dev* and *fmg* transcripts in a *ladA* mutant. Wild-type DK1622 and its *ladA* mutant derivative were subjected to starvation under submerged culture conditions and samples were collected at the indicated number of hours poststarvation (PS) for measurement of (A) *dev*, (B) *fmgA*, (C) *fmgBC*, (D) *fmgD*, and (E) *fmgE* transcript levels by RT-qPCR. Graphs show the data points and average of three biological replicates, relative to wild-type DK1622 at 18 h PS, and error bars show one standard deviation. Asterisks indicate a significant difference ($p < 0.05$ in Student's two-tailed $t$-tests) from wild type at the corresponding time PS.

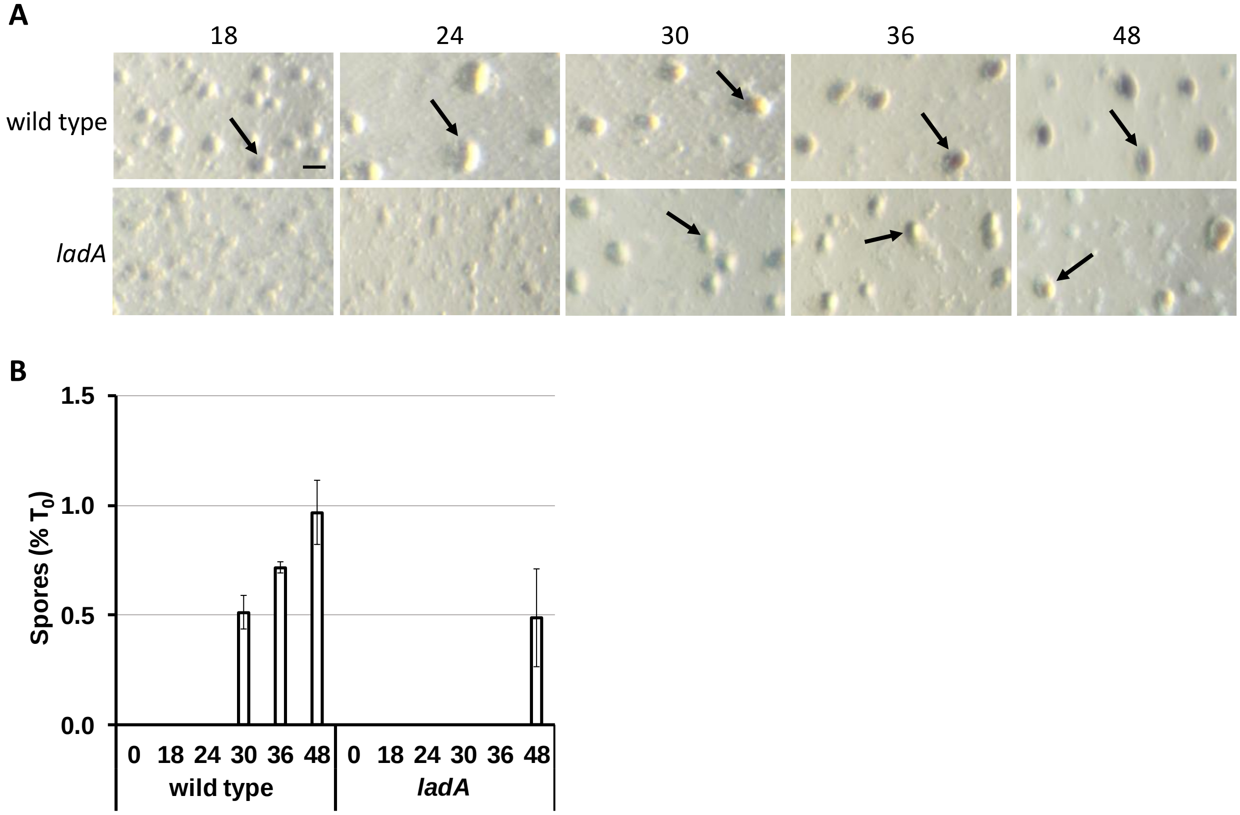


Figure S6. Development of the *ladA* mutant. Wild-type DK1622 and its *ladA* mutant derivative were subjected to starvation under submerged culture conditions. (A) Microscopy. Images were obtained at the indicated number of hours poststarvation (PS). DK1622 formed mounds by 18 h PS (an arrow points to one) and the mounds darkened at 36 to 48 h. The *ladA* mutant formed mounds at 30 h, and the mounds did not darken until 48 h. Bar, 100 mm. Similar results were observed in at least three biological replicates. (B) Quantification of sonication-resistant spores. Values are expressed as a percentage of the number of rod-shaped cells present at the time starvation initiated development ($T_0$) (Table S1). Bars show the average of three biological replicates and error bars show one standard deviation.

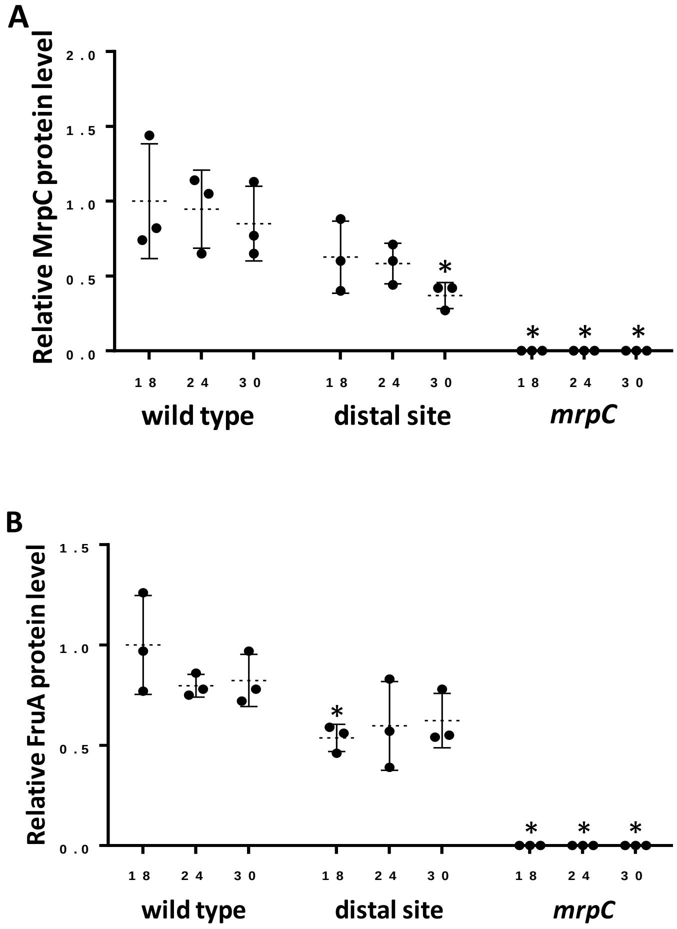


Figure S7. Levels of MrpC and FruA during *M. xanthus* development. Wild-type DK1622 and its indicated mutant derivatives were subjected to starvation under submerged culture conditions and samples were collected at the indicated number of hours poststarvation (PS) for measurement of (A) MrpC and (B) FruA levels by immunoblot. Graphs show the data points and average of at least three biological replicates, relative to wild-type DK1622 at 18 h PS, and error bars show one standard deviation. Asterisks indicate a significant difference ($p < 0.05$ in Student's two-tailed *t*-tests) from wild type at the corresponding time PS.

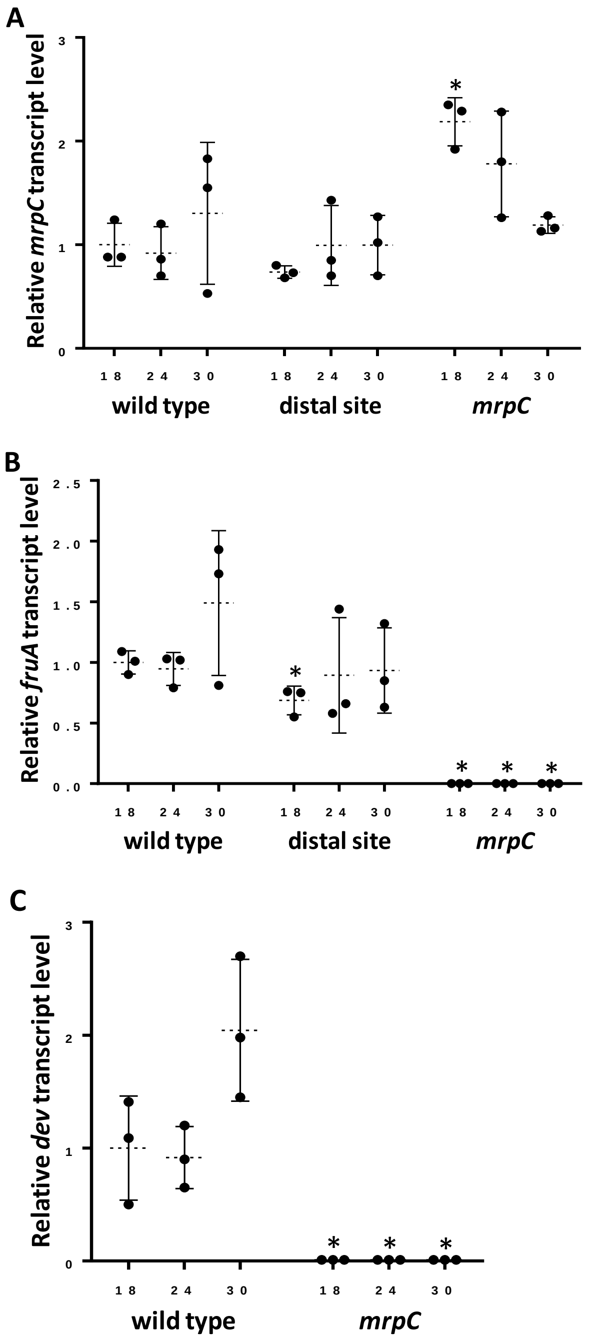


Figure S8. Transcript levels during *M. xanthus* development. Wild-type DK1622 and its indicated mutant derivatives were subjected to starvation under submerged culture conditions and samples were collected at the indicated number of hours poststarvation (PS) for measurement of (A) *mrpC*, (B) *fruA*, and (C) *dev* transcript levels by RT-qPCR. Graphs show the data points and average of at least three biological replicates, relative to wild-type DK1622 at 18 h PS, and error bars show one standard deviation. Asterisks indicate a significant difference ($p$ < 0.05 in Student's two-tailed $t$-tests) from wild type at the corresponding time PS.

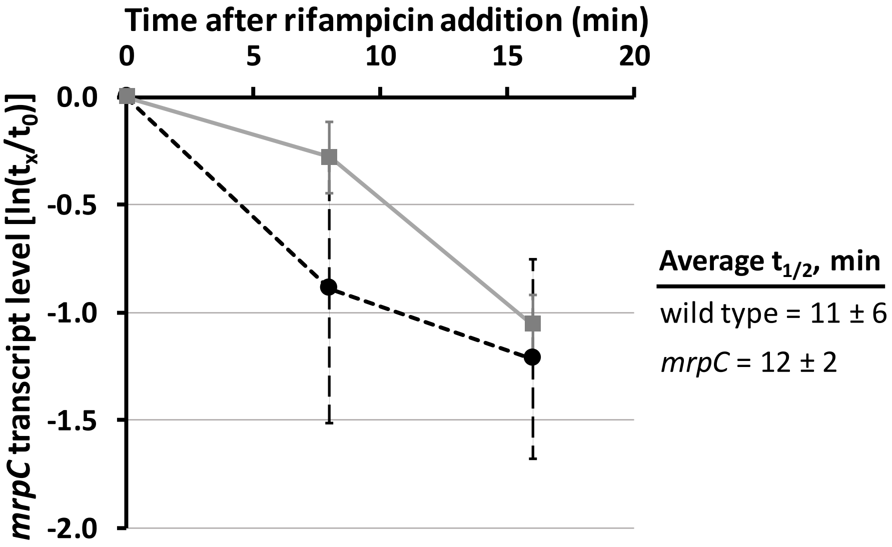


Figure S9. *mrpC* transcript stability. Wild-type DK1622 and the *mrpC* mutant were subjected to starvation under submerged culture conditions for 18 h. The overlay was replaced with fresh starvation buffer containing rifampicin (50 μg/ml) and samples were collected immediately ($t_0$) and at the times indicated ($t_x$) for measurement of the *mrpC* transcript level by RT-qPCR. Transcript levels at $t_x$ were normalized to that at $t_0$ for each of three biological replicates and used to determine the transcript half-life for each replicate. The average half-life (Average $t_{1/2}$) and one standard deviation are shown, and the difference is not statistically significant ($p$ = 0.85 in a Student's two-tailed *t*-test). The graph shows the average $\ln(t_x/ t_0)$ and one standard deviation for the three biological replicates of wild type (black dashed line) and the *mrpC* mutant (gray solid line).

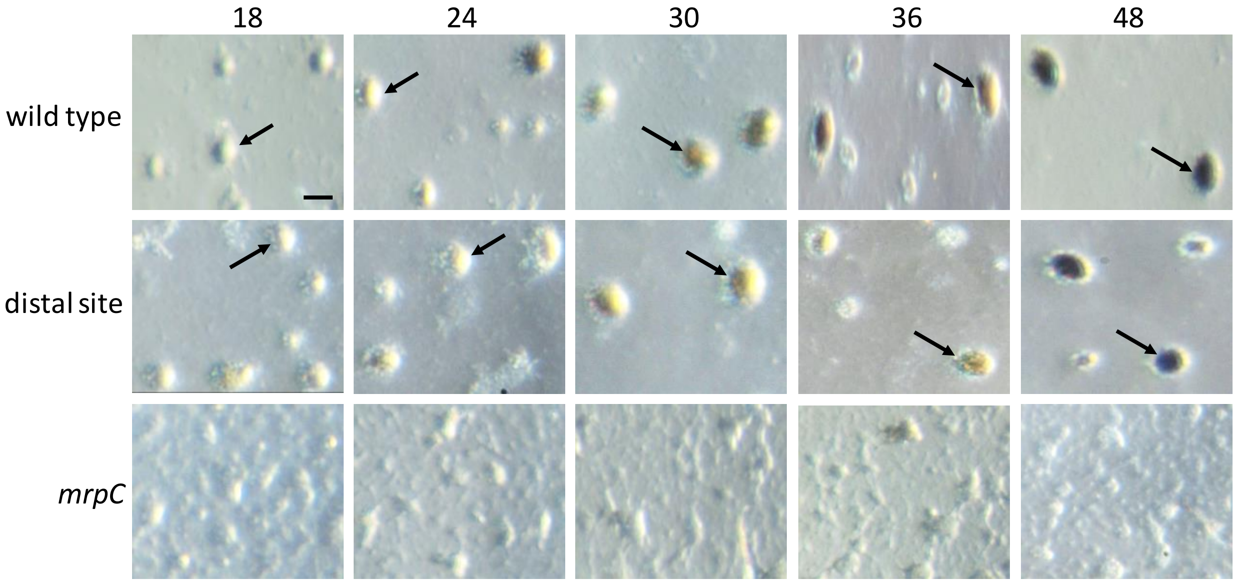


Figure S10. Development of *M. xanthus* strains. Wild-type DK1622 and its indicated mutant derivatives were subjected to starvation under submerged culture conditions and images were obtained at the indicated number of hours poststarvation (PS). The wild type and the distal site mutant formed mounds by 18 h PS (an arrow points to one) and the mounds darkened at 36 to 48 h. The *mrpC* mutant failed to form mounds. Bar, 100 μm. Similar results were observed in at least three biological replicates.

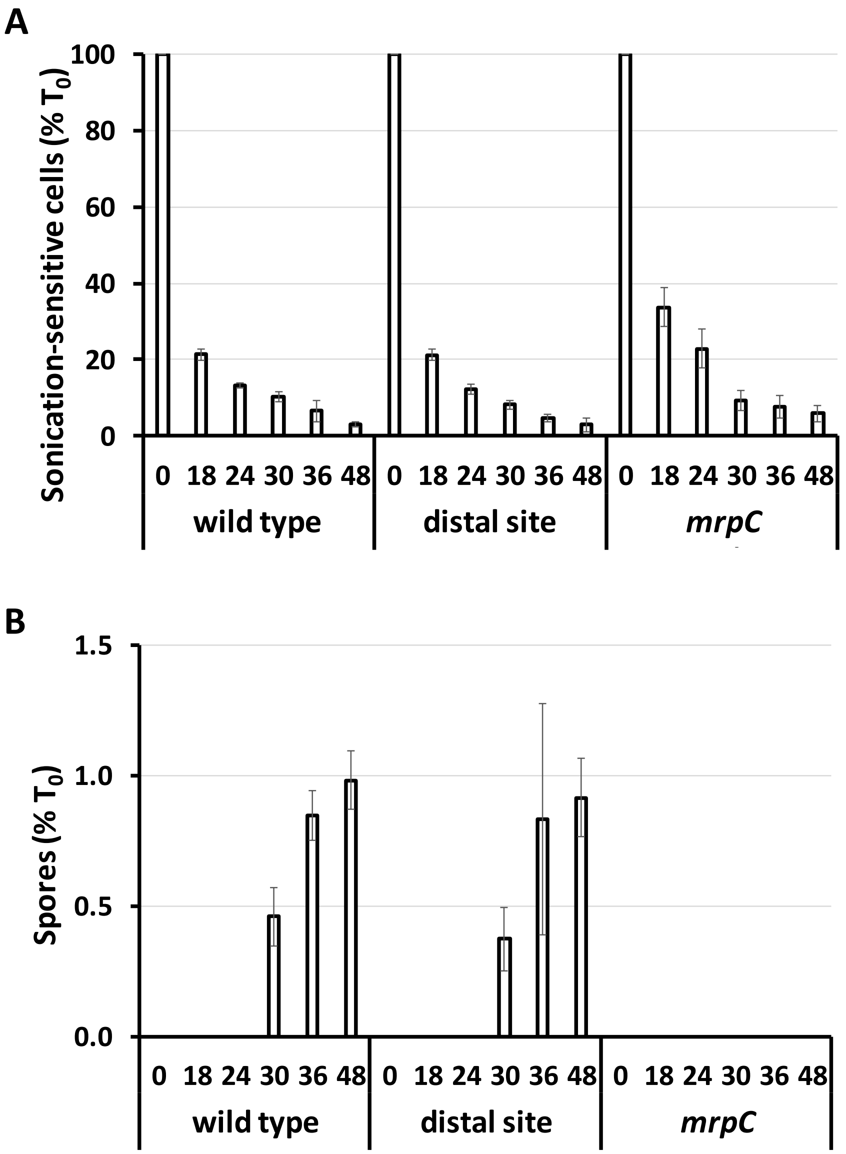


Figure S11. Cellular changes during *M. xanthus* development. Wild-type DK1622 and its indicated mutant derivatives were subjected to starvation under submerged culture conditions and samples were collected at the indicated number of hours poststarvation for quantification of (A) sonication-sensitive cells and (B) sonication-resistant spores. Values are expressed as a percentage of the number of rod-shaped cells present at the time starvation initiated development ($T_0$) (Table S1). Bars show the average of three biological replicates and error bars show one standard deviation.

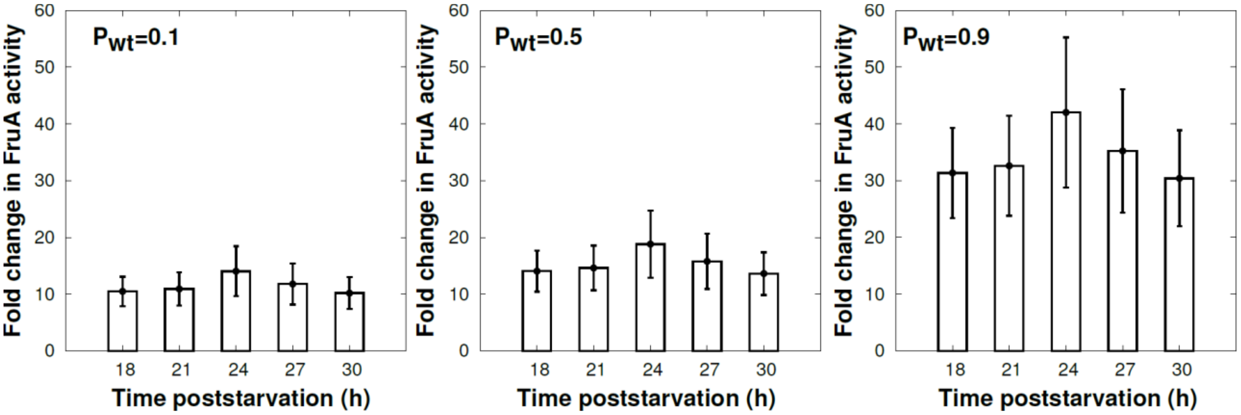


Figure S12. Mathematical modeling prediction of the required reduction in FruA activity in the *csgA* mutant in comparison to wild type, to explain the experimental data. Plots are for different values of promoter saturation ($P_{wt}$) at the indicated times poststarvation. Bars show the average of 108 datasets representing all possible combinations of four biological replicates of wild type and three biological replicates of each mutant (*csgA*, *devI*, *devS*), and error bars show one standard deviation.

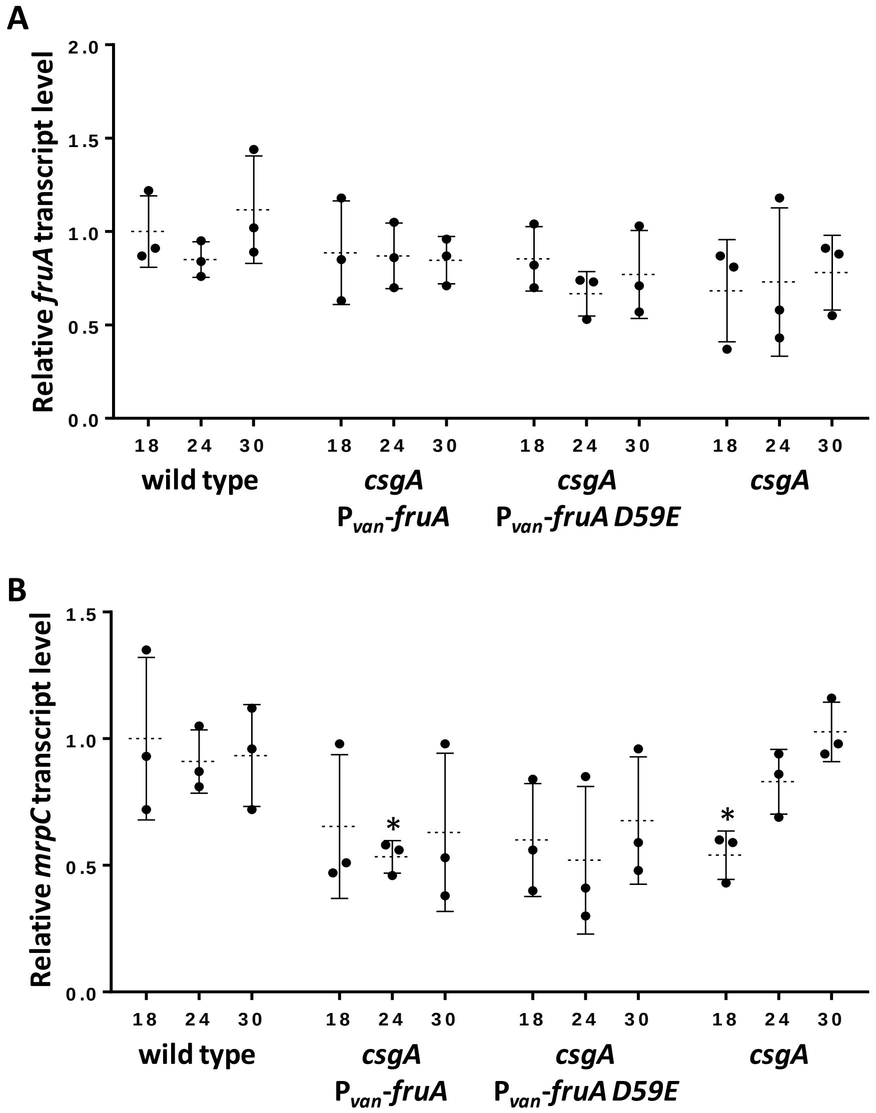


Figure S13. Transcript levels during *M. xanthus* development. Wild-type DK1622 and its indicated mutant derivatives were subjected to starvation under submerged culture conditions and samples were collected at the indicated number of hours poststarvation (PS) for measurement of (A) *fruA* and (B) *mrpC* transcript levels by RT-qPCR. Bars show the average of at least three biological replicates, relative to wild-type DK1622 at 18 h PS, and error bars show one standard deviation. Asterisks indicate a significant difference ($p$ < 0.05 in Student's two-tailed $t$-tests) from wild type at the corresponding time PS.

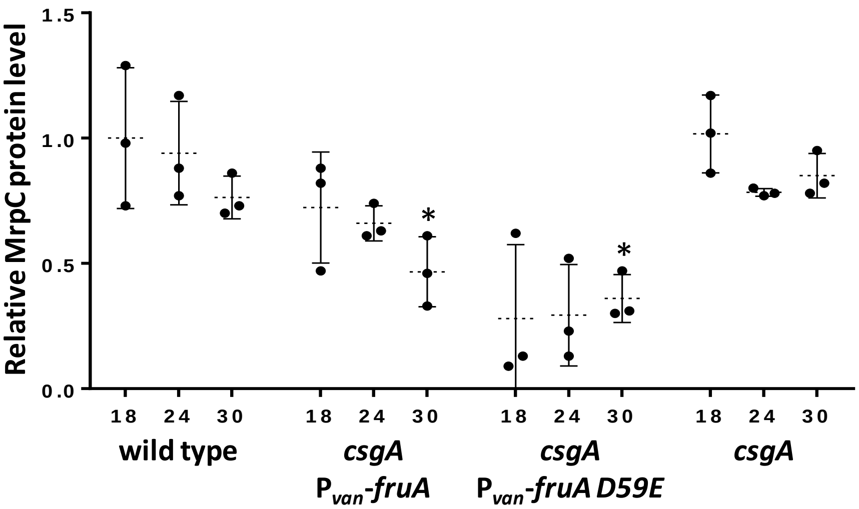


Figure S14. Level of MrpC during *M. xanthus* development. Wild-type DK1622 and its indicated mutant derivatives were subjected to starvation under submerged culture conditions and samples were collected at the indicated number of hours poststarvation (PS) for measurement of MrpC levels by immunoblot. Bars show the average of at least three biological replicates, relative to wild-type DK1622 at 18 h PS, and error bars show one standard deviation. Asterisks indicate a significant difference ($p < 0.05$ in Student's two-tailed $t$-tests) from wild type at the corresponding time PS.

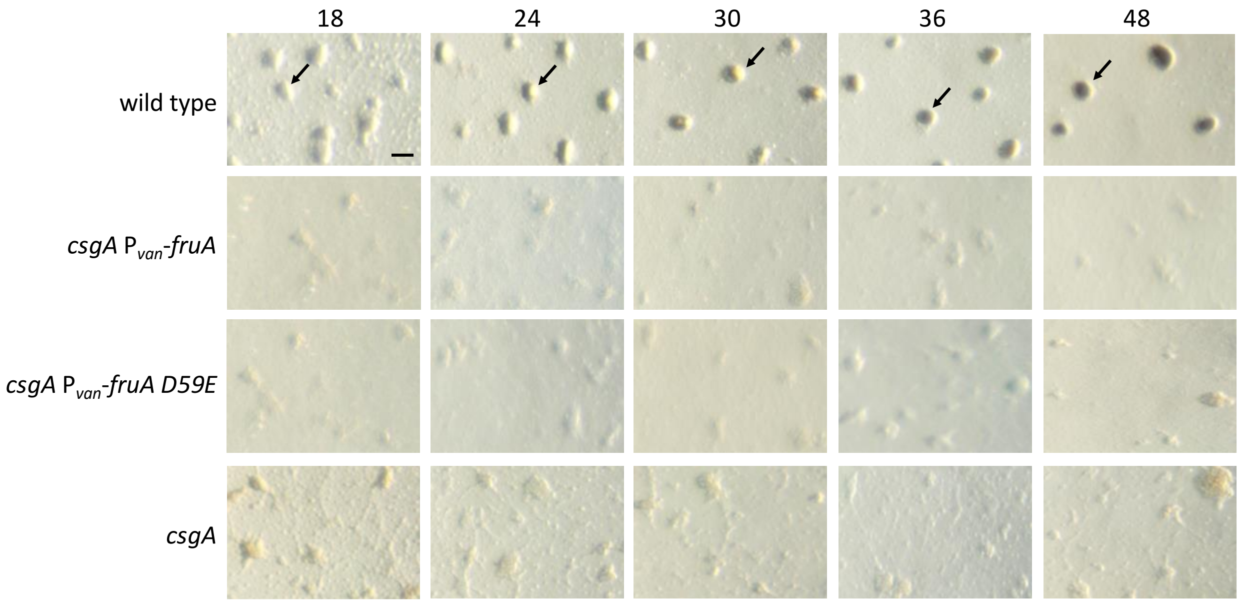


Figure S15. Development of *M. xanthus* strains. Wild-type DK1622 and its indicated mutant derivatives were subjected to starvation under submerged culture conditions and images were obtained at the indicated number of hours poststarvation (PS). The wild type formed mounds by 18 h PS (an arrow points to one) and the mounds darkened at 36 to 48 h. The *csgA* P$_{van}$-*fruA*, *csgA* P$_{van}$-*fruA D59E*, and *csgA* mutants failed to form mounds. Bar, 100 mm. Similar results were observed in at least three biological replicates.

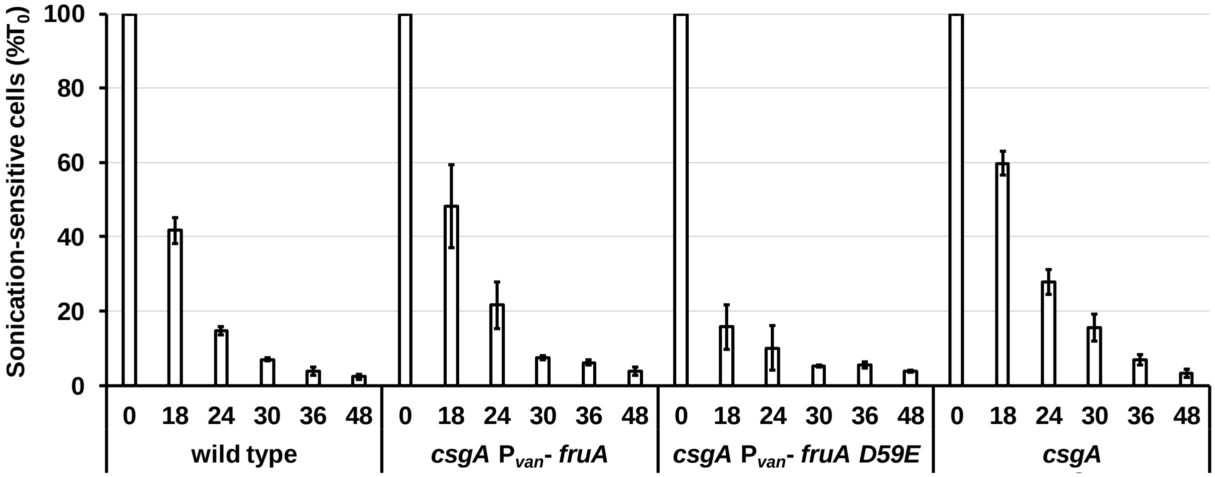


Figure S16. Cellular changes during *M. xanthus* development. Wild-type DK1622 and its indicated mutant derivatives were subjected to starvation under submerged culture conditions and samples were collected at the indicated number of hours poststarvation for quantification of sonication-sensitive cells. Values are expressed as a percentage of the number of rod-shaped cells present at the time starvation initiated development ($T_0$) (Table S1). Bars show the average of three biological replicates and error bars show one standard deviation.

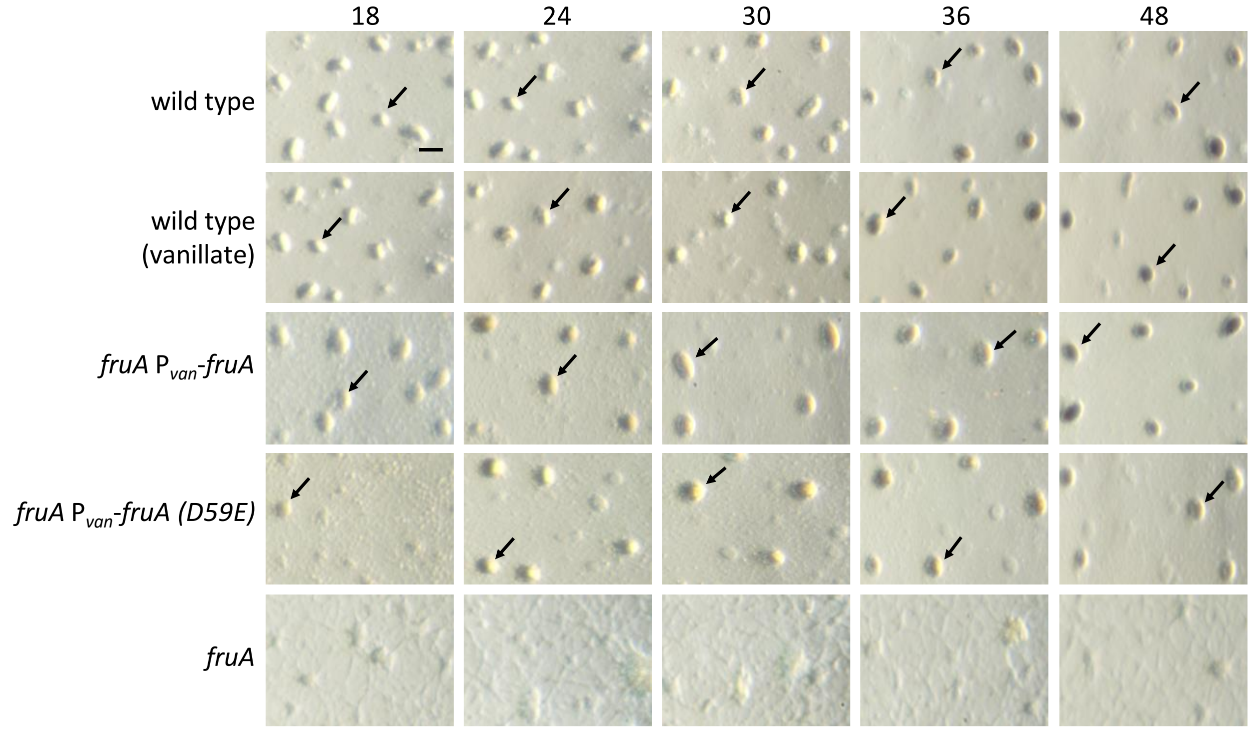


Figure S17. Development of *M. xanthus* strains. Wild-type DK1622 without or with vanillate induction and its indicated mutant derivatives with vanillate induction were subjected to starvation under submerged culture conditions and images were obtained at the indicated number of hours poststarvation (PS). The wild type without or with vanillate, and the *fruA* $P_{van}$-*fruA* and *fruA* $P_{van}$-*fruA D59E* strains, formed mounds by 18 h PS (arrows point to mounds) and the mounds darkened at 36 to 48 h. The *fruA* mutant failed to form mounds. Bar, 100 mm. Similar results were observed in at least three biological replicates.

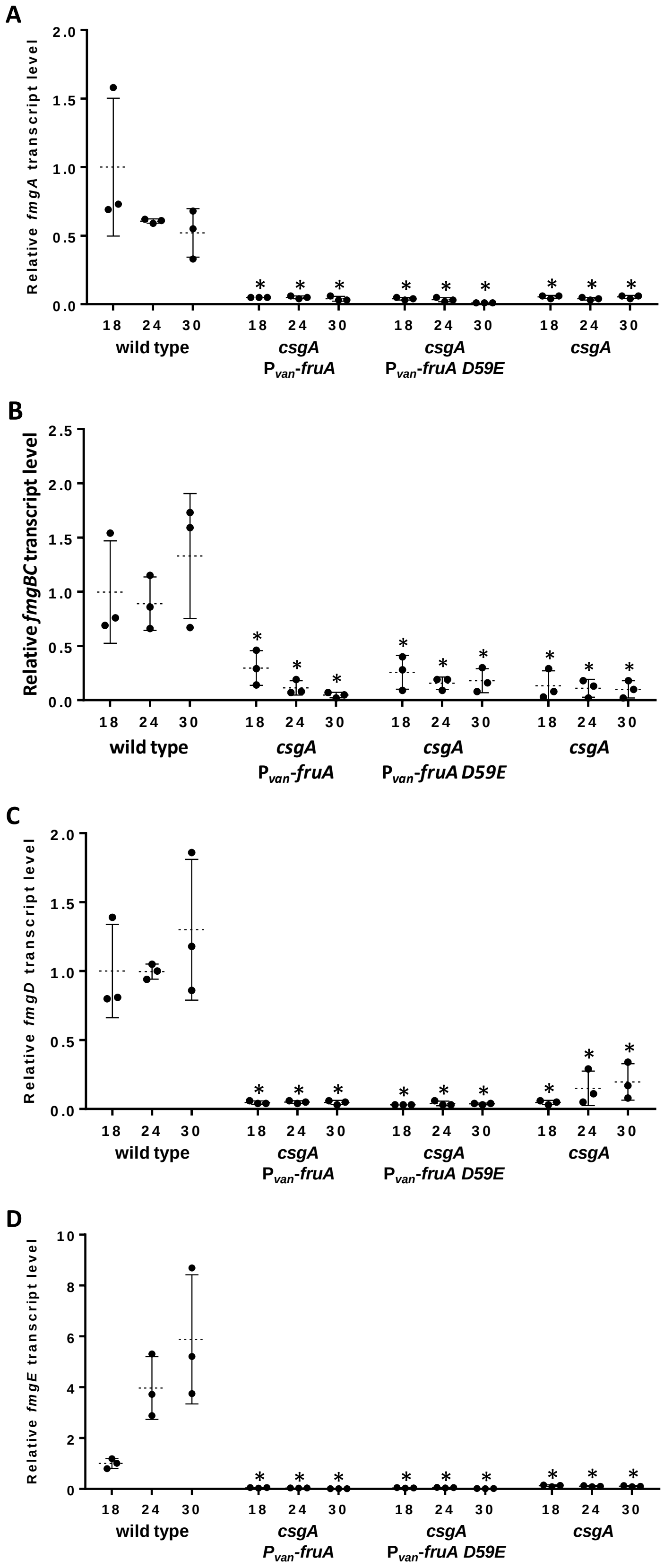
A
Relative fmgA transcript level
2.0
1.5
1.0
0.5
0.0
18 24 30
wild type
csgA
Pvan-fruA
csgA
Pvan-fruA D59E
csgA
B
Relative fmgBC transcript level
2.5
2.0
1.5
1.0
0.5
0.0
C
Relative fmgD transcript level
D
Relative fmgE transcript level
10
8
6
4
2
0

Figure S18. Levels of *fmg* transcripts in *csgA* mutants. Wild-type DK1622 and its indicated mutant derivatives were subjected to starvation under submerged culture conditions and samples were collected at the indicated number of hours poststarvation (PS) for measurement of (A) *fmgA*, (B) *fmgBC*, (C) *fmgD*, and (D) *fmgE* transcript levels by RT-qPCR. Bars show the average of at least three biological replicates, relative to wild-type DK1622 at 18 h PS, and error bars show one standard deviation. Asterisks indicate a significant difference ($p < 0.05$ in Student's two-tailed *t*-tests) from wild type at the corresponding time PS.

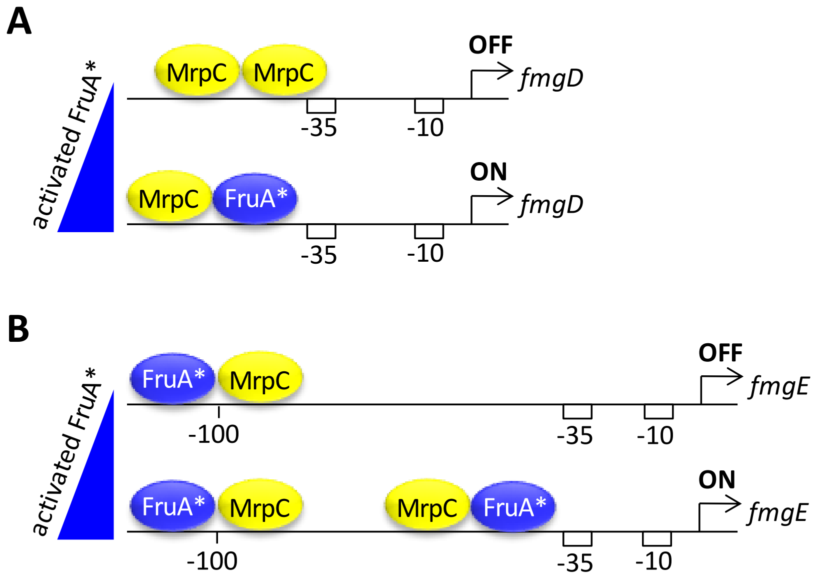


Figure S19. Models for regulation of *fmgD* and *fmgE*. C-signaling causes the level of activated FruA* to rise as development proceeds (triangles). (A) Cooperative binding of two MrpC initially represses *fmgD* transcription, but eventually FruA* outcompetes the downstream MrpC for binding to the upstream MrpC, activating transcription. (B) MrpC and activated FruA* bind cooperatively first to a higher affinity centered at -100 bp relative to the *fmgE* transcriptional start site. As FruA* rises, the lower affinity site just upstream of the promoter is also cooperatively bound by FruA* and MrpC, activating transcription. In both panels, boxes indicate the promoter -35 and -10 regions.

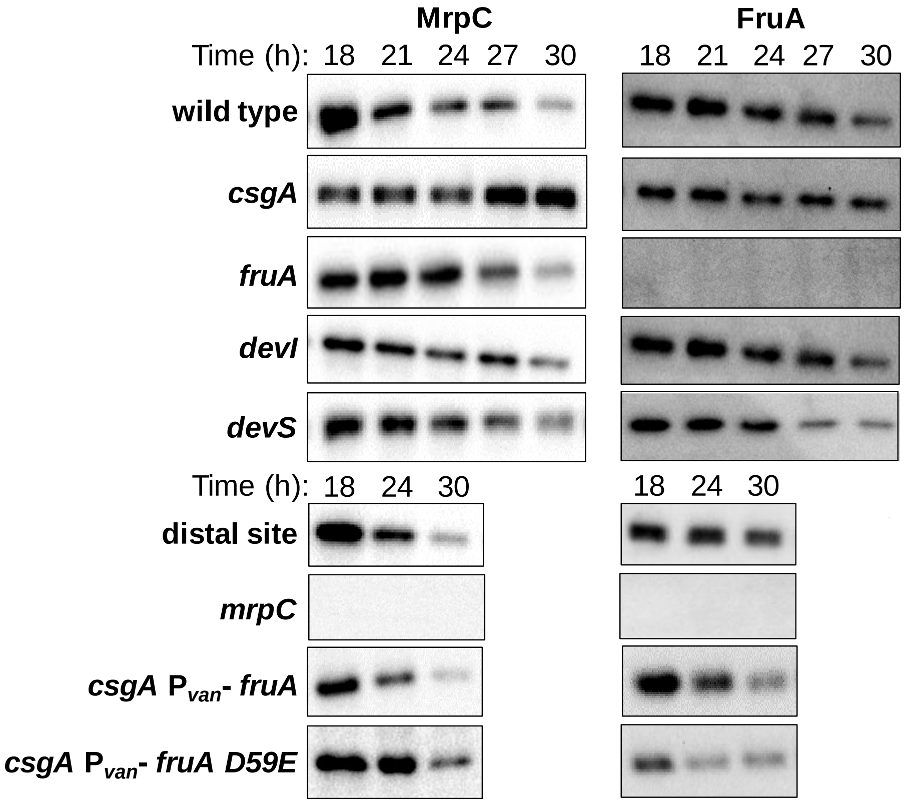


Figure S20. Representative MrpC and FruA immunoblots for wild type and mutants. Wild-type DK1622 and its indicated mutant derivatives were subjected to starvation under submerged culture conditions and samples were collected at the indicated number of hours poststarvation (PS) for measurement of MrpC and FruA levels by immunoblot. Equal volumes (10 μl for measurement of MrpC and 15 μl for measurement of FruA) of whole-cell extract samples were subjected to semi-quantitative immunoblot analysis as described in the Experimental Procedures.

### References


Campbell, A., P. Viswanathan, T. Barrett, B. Son, S. Saha & L. Kroos, (2015) Combinatorial regulation of the *dev* operon by MrpC2 and FruA during *Myxococcus xanthus* development. *J. Bacteriol.* **197**: 240-251.

Hanahan, D., (1983) Studies on transformation of *Escherichia coli* with plasmids. *J. Mol. Biol.* **166**: 557-580.

Iniesta, A. A., F. Garcia-Heras, J. Abellon-Ruiz, A. Gallego-Garcia & M. Elias-Arnanz, (2012) Two systems for conditional gene expression in *Myxococcus xanthus* inducible by isopropyl-β-D-thiogalactopyranoside or vanillate. *J. Bacteriol.* **194**: 5875-5885.

Kaiser, D., (1979) Social gliding is correlated with the presence of pili in *Myxococcus xanthus*. *Proc. Natl. Acad. Sci. USA* **76**: 5952-5956.

Kroos, L. & D. Kaiser, (1987) Expression of many developmentally regulated genes in *Myxococcus* depends on a sequence of cell interactions. *Genes Dev.* **1**: 840-854.

Ossa, F., M. E. Diodati, N. B. Caberoy, K. M. Giglio, M. Edmonds, M. Singer & A. G. Garza, (2007) The *Myxococcus xanthus* Nla4 protein is important for expression of stringent response-associated genes, ppGpp accumulation, and fruiting body development. *J. Bacteriol.* **189**: 8474-8483.

Rajagopalan, R. & L. Kroos, (2014) Nutrient-regulated proteolysis of MrpC halts expression of genes important for commitment to sporulation during *Myxococcus xanthus* development. *J. Bacteriol.* **196**: 2736-2747.

Rajagopalan, R. & L. Kroos, (2017) The *dev* operon regulates the timing of sporulation during *Myxococcus xanthus* development. *J. Bacteriol.* **199**: e00788-00716.

Rajagopalan, R., S. Wielgoss, G. Lippert, G. J. Velicer & L. Kroos, (2015) *devI* is an evolutionarily young negative regulator of *Myxococcus xanthus* development. *J Bacteriol* **197**: 1249-1262.

Shimkets, L. J. & S. J. Asher, (1988) Use of recombination techniques to examine the structure of the *csg* locus of *Myxococcus xanthus*. *Mol. Gen. Genet.* **211**: 63-71.

Sun, H. & W. Shi, (2001) Genetic studies of *mrp*, a locus essential for cellular aggregation and sporulation of *Myxococcus xanthus*. *J. Bacteriol.* **183**: 4786-4795.

Viswanathan, P., K. Murphy, B. Julien, A. G. Garza & L. Kroos, (2007a) Regulation of *dev*, an operon that includes genes essential for *Myxococcus xanthus* development and CRISPR-associated genes and repeats. *J. Bacteriol.* **189**: 3738-3750.

Viswanathan, P., T. Ueki, S. Inouye & L. Kroos, (2007b) Combinatorial regulation of genes essential for *Myxococcus xanthus* development involves a response regulator and a LysR-type regulator. *Proc. Natl. Acad. Sci. USA* **104**: 7969-7974.
